# Glycated alpha-synuclein assemblies cause distinct Parkinson’s disease pathogenesis in mice

**DOI:** 10.1101/2025.06.04.657879

**Authors:** Akshaya Rajan, Anish Varghese, Shaliya Puthanveedu Hashardeen, Ann Teres Babu, Vinesh Vijayan, Poonam Thakur

**Affiliations:** School of Biology, IISER-Thiruvananthapuram, Kerala, India-695551; Institute of Structural Biology, Rudolf Virchow Center for Integrative and Translational Bioimaging, Julius-Maximilians-Universität Würzburg, Josef-Schneider-Straße 2, 97080 Würzburg, Germany; School of Chemistry, IISER-Thiruvananthapuram, Kerala, India-695551

**Keywords:** Advanced Glycation End products, Receptor for Advanced Glycation End products, neuroinflammation, phospho-synuclein, hyperglycemia, diabetes, Substantia Nigra

## Abstract

Alpha-synuclein (α-Syn) misfolding and aggregation are key drivers of Parkinson’s disease (PD) pathology. Mutations and certain post-translational modifications impact its aggregation propensity and pathogenicity. Glycation, a non-enzymatic modification enhanced during hyperglycemia and aging, both known risk factors for PD, has been implicated in α-Syn pathology. Although preformed α-Syn-fibrils induce PD-like phenotypes in mice, the impact of glycation on their pathogenicity is unclear. In the current study, we glycated α-Syn using methylglyoxal (MGO), a potent glycating agent, resulting in altered biophysical characteristics in comparison to non-glycated α-Syn. Glycation inhibited the formation of typical beta sheet structures under aggregating conditions. Despite that, glycated α-Syn assemblies induced dopaminergic neurodegeneration and neuroinflammation to a similar extent as the non-glycated α-Syn fibrils upon their injection in the mouse substantia nigra (SN). However, these glycated assemblies triggered higher neuroinflammation and increased accumulation of receptors for advanced glycation end products (RAGE) compared to non-glycated fibrils. Consequently, an earlier onset of neuromuscular deficits and anxiety was observed in these mice. Thus, glycation of α-Syn causes distinct PD-associated pathology compared to non-glycated α-Syn, causing an earlier onset of motor symptoms. These findings provide insight into how glycation of α-Syn due to hyperglycemia may contribute to an increased risk of PD in diabetic populations.

## 1. INTRODUCTION

Parkinson’s disease (PD) is a progressive neurodegenerative disorder affecting approximately 2-3% of the global population aged over 60 [1,2]. One of the major pathological hallmarks of PD is the misfolding and accumulation of alpha-synuclein (α-Syn) into Lewy bodies [3,4]. This causes dopaminergic neurodegeneration in the substantia nigra (SN) region of the brain [5,6], resulting in motor symptoms characteristic of PD [7,8]. Although α-Syn is intrinsically disordered protein, its structure is influenced by molecular interactions and post-translational modifications, which can disrupt homeostasis and drive misfolding during disease progression [9,10]. Among different post-translational modifications, glycation, a non-enzymatic modification, has emerged as a key factor promoting the conversion of α-Syn into pathological species [11]. Glycation generates irreversible advanced glycation end products (AGEs), which are known to accumulate at the periphery of Lewy bodies and are reported in post-mortem PD brain samples [12,13]. The likelihood of glycation and subsequent AGE accumulation increases substantially during metabolic disorders, particularly in the hyperglycemic conditions associated with type-2 diabetes mellitus [14]. Importantly, multiple epidemiological studies have demonstrated an elevated risk of developing PD in diabetic patients, even when hyperglycemia is effectively managed using standard anti-diabetic medications [15,16]. This has led to the hypothesis that fluctuations in glucose levels, rather than sustained hyperglycemia per se, may exacerbate diabetic complications and increase glycation risks for biomolecules such as α-Syn [17]. Such glucose fluctuations are common in both pre-diabetic and diabetic people. While diabetes has also been linked to other neurological disorders such as Alzheimer’s disease [18] and peripheral neuropathy [19], the specific molecular mechanisms that predispose some diabetic individuals to PD remain poorly understood.

It is hypothesized that the glycation of α-Syn may represent a critical molecular link in this increased PD risk. Some studies in the literature have begun to address this possibility. α-Syn upon glycation has been shown to alter its aggregation kinetics and fibril formation capabilities *in vitro* [11,20]. At the functional level, glycated α-Syn exhibits impaired membrane binding and interferes with normal ubiquitination and clearance processes [21,22]. The altered aggregation propensity also disrupts its physiological ability to bind and fuse with synaptic vesicles, thereby potentially contributing to synaptic dysfunction and neurodegeneration [23]. In α-Syn-overexpressing mouse models of PD, long-term hyperglycemia induced by streptozotocin exacerbates motor deficits, α-Syn aggregation, neuroinflammation, and the loss of tyrosine hydroxylase (TH)-positive neurons in the SN [24]. Similarly, direct infusion of the glycating agent methylglyoxal (MGO) into the SN of mice promotes α-Syn accumulation, AGE deposition, and dopaminergic neurodegeneration [22]. MGO-mediated glycation of α-Syn not only enhances its aggregation propensity but also potentiates neuroinflammatory responses, suggesting a mechanism by which altered proteostasis may contribute to the increased risk of developing PD in individuals with diabetes [12]. Supporting this, intracerebroventricular infusion of MGO in α-Syn transgenic mice also results in PD behavioral phenotypes, alongside increased α-Syn and AGE accumulation in the midbrain [25]. These findings strongly indicate that a glycating environment can profoundly dysregulate the dopaminergic system.

While these important studies have clearly demonstrated the deleterious effects of a generalized glycating environment in the brain, several unanswered questions remain. Specifically, it is unclear whether the overall glycating milieu enhances PD risk through multiple mechanisms or whether glycated α-Syn alone is sufficient to exacerbate PD pathology compared to non-glycated fibrils. Recent findings suggest that glycated α-Syn may trigger a more specific pathological cascade, including activation of the TLR2 and NLRP3 inflammasome pathways, leading to robust microglial cell activation *in vitro* and promoting dopaminergic neurodegeneration *in vivo* [26]. Additionally, glycated α-Syn has been shown to significantly enhance the expression of the receptor for advanced glycation end products (RAGE) when stereotaxically injected into the SN of mice [27], suggesting a potential feed-forward mechanism of neuroinflammation and neurodegeneration.

In the current study, we directly investigate the impact of glycated α-Syn on dopaminergic neurodegeneration in mice and systematically compare its effects with non-glycated α-Syn fibrils to determine whether glycation specifically exacerbates PD pathology. While most published studies have focused exclusively on male mice, we have included both male and female mice in our experimental design to provide a more comprehensive perspective on sex differences. Our findings demonstrate that while glycation inhibits the formation of long, mature α-Syn fibrils, glycated α-Syn assemblies still induce similar levels of neurodegeneration as the non-glycated α-Syn fibrils. Interestingly, brains injected with glycated α-Syn show lower accumulation of phosphorylated α-Syn (p-Syn) but exhibit significantly higher levels of RAGEs. These elevated RAGE levels could further exacerbate neuroinflammation and, in turn, accelerate behavioral deficits. Together, these results suggest that glycated α-Syn drives neurodegeneration through mechanisms distinct from those of non-glycated fibrils, providing new insights into how diabetes may increase PD risk through protein glycation.

## 2. RESULTS AND DISCUSSION

### 2.1. MGO inhibits α-Syn beta sheet fibril formation under aggregating conditions

The potential modifying effects of MGO on α-Syn aggregation were investigated by incubating purified α-Syn monomers in the presence of MGO for 96 hours at 37°C with shaking (Figure 1A). Unmodified fibrils were generated by incubating α-Syn monomers in a similar volume of 1X DPBS (Dulbecco’s phosphate-buffered saline). The resulting assemblies were termed MGO-F (with MGO) and DPBS-F (without MGO). Thioflavin T (ThT) fluorescence assay revealed that DPBS-F exhibited a characteristic rise (∼60-fold increase) compared to MGO-F in ThT fluorescence, consistent with cross beta-sheet structure formation (Figure 1B). MGO markedly hindered this fibril formation, as indicated by the minimal increase in ThT fluorescence over time. In contrast, the monomeric control, which was not subjected to aggregating conditions, showed a basal level of ThT fluorescence. Circular dichroism (CD) spectroscopy was used to further validate these findings (Figure 1C). While DPBS-F displayed an ellipticity minima around 220 nm, typical of beta sheet structures, MGO-F retained a random coil conformation similar to the monomer. This suggests that in aggregating conditions, the structural transition of α-Syn from a random coil to a beta sheet-rich fibrillary structure, a hallmark of amyloid formation, is prevented by the presence of MGO.

**Figure 1.**
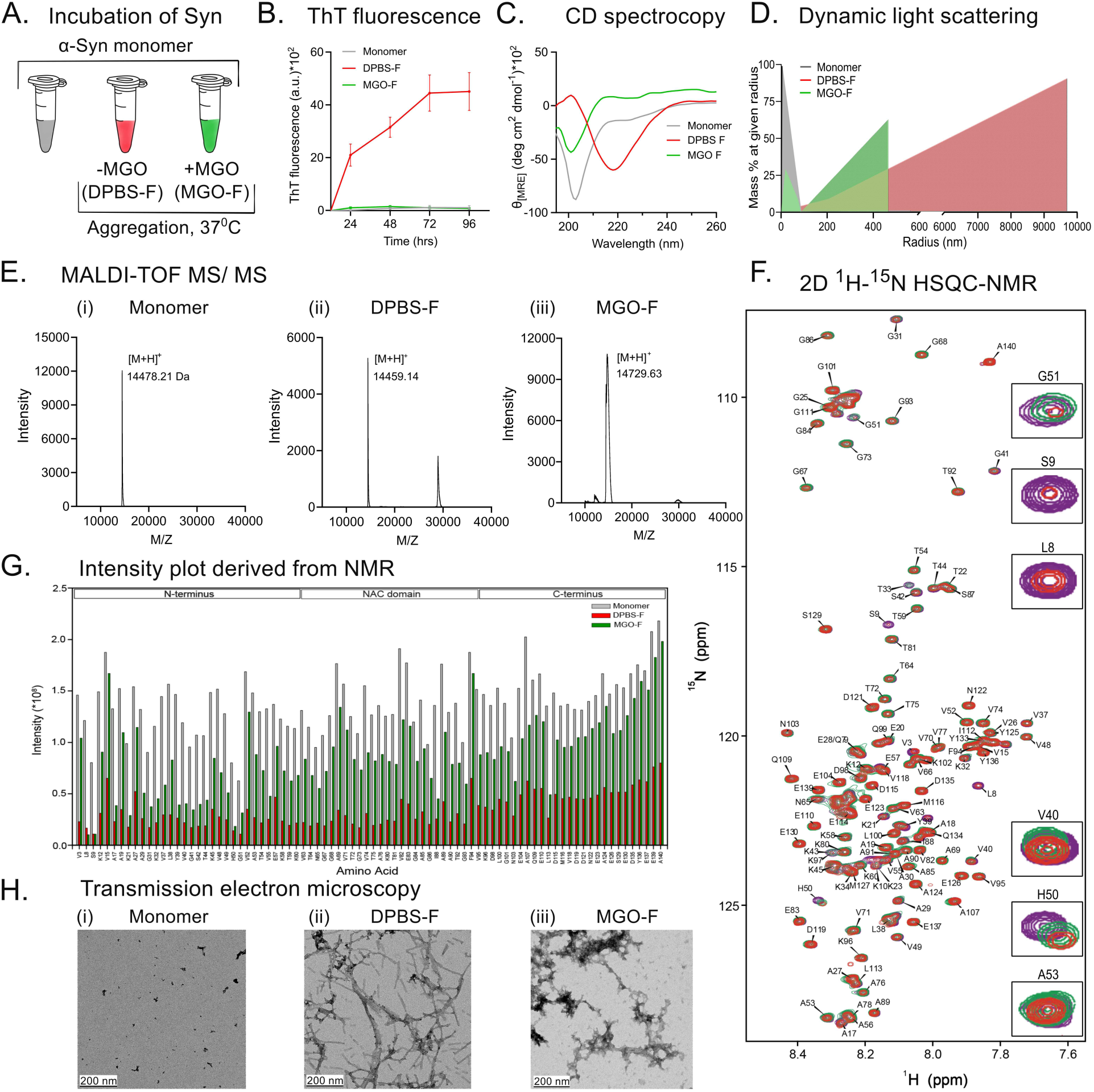
- Glycated α-Syn forms amorphous assemblies, distinct from amyloid fibril structure. A. Purified recombinant α-Syn monomer was incubated at 37°C with shaking in the presence (MGO-F) or absence (DPBS-F) of MGO, to induce aggregation B. Average ThT fluorescence of α-Syn species monitored throughout the incubation phase shows accumulating cross beta-sheet structure in DPBS-F compared to MGO-F, mean ± SEM, n=3. C. Representative CD spectroscopy of α-Syn monomer control and α-Syn assemblies with and without MGO after 96 hours of incubation confirms the absence of beta-sheet structure in MGO-F and monomer. D. Representative DLS histogram showing a heterogeneous particle range in MGO-F in contrast to large-sized particles in DPBS-F. E. Representative MALDI-TOF spectra of α-Syn monomer (i) and α-Syn incubated without (ii) and with MGO (iii) for 96 hours, indicating an increase in mass of α-Syn with MGO incubation. F. Overlay of 2D ^1^H-^15^N HSQC NMR spectra of recombinant ^15^N labeled α-Syn without (red) and with (green) MGO and α-Syn monomeric control (purple), inset shows zoomed in images of selected amino acid residues displaying contour peak differences. G. Absolute intensity plots of DPBS-F (red) and MGO-F (green) compared with monomer (grey) show lower intensities of DPBS-F. H. Representative TEM images of monomeric α-Syn control (i), and α-Syn incubated without and with MGO (ii and iii) for 96 hours show the absence of long, fibrillar structures in MGO-F.

Dynamic light scattering (DLS) analysis revealed distinct differences in the size distribution of these assemblies (Figure 1D). Monomers exhibited a narrow size distribution with a predominant radius of 3 nm, while DPBS-F formed larger particles mostly possessing radii of about 9700 nm, consistent with the presence of long fibrillar assemblies. In contrast, MGO-F displayed a heterogeneous population of particles with radii ranging between those of monomers and fibrils. This heterogeneity likely arises from the random nature of glycation, which generates diverse AGEs with varying molecular masses and abundance of lysine residues in α-Syn, most of which can be modified by MGO [28]. Interestingly, 11 of the total 15 lysines are located in its N-terminus half and this region of α-Syn is known to influence its aggregation status [29]. Thus, the modified N-terminal lysines might be contributing to hampered interactions between the amino acid residues and preventing the formation of long fibrils. MALDI-TOF mass spectrometry analysis indeed shows that MGO-F displayed an additional mass of ∼252 Da compared to the monomer (Figure 1E). This mass shift corresponds with the incorporation of MGO residues into α-Syn. The broader peak of MGO-F also explains a composition with the various size-ranges in MGO-F compared to the sharp peaks in the case of monomer and DPBS-F. The appearance of the second peak in DPBS-F indicates the presence of higher molecular weight species, which are barely discernible in MGO-F and absent in monomers.

To further characterize the impact of glycation on α-Syn, we performed 2D ^1^H-^15^N HSQC NMR spectroscopy on ^15^N-labeled α-Syn (Figure 1F). With respect to NMR spectra, the contour peak differences were observed in only a few of the lysine residues upon glycation. Generally, highly soluble intrinsically disordered proteins will have high protein dynamics, resulting in relatively sharp peaks (high intensity) in NMR. However, as the protein aggregates, the protein dynamics slow down, and as a result, the NMR peaks broadens causing the NMR peak intensities to drop. After 96 hours of incubation, DPBS-F showed a significant reduction in peak intensities, indicative of fibril formation (Figure 1G). In contrast, MGO-F retained higher peak intensities, suggesting that MGO reduces the aggregation rate and prevents the formation of large, NMR-invisible fibrils. However, the peak intensities of MGO-F were still lower than the non-incubated monomer, especially at the Lys-rich N-terminal region. This confirms that in the presence of MGO, even in the aggregating conditions, α-Syn will not form aggregates but leads to an assembly distinct from monomers. In addition, we observed a smaller reduction in intensities upon glycation for amino acids contributing to hydrophobicity, compared to the non-glycated sample and the monomeric control. So, the glycation modification on lysine might have indirectly affected the hydrophobicity of the protein, preventing its aggregation. The structural polymorphism of α-Syn has been extensively characterized, and fibrils isolated from the brains of PD patients have been shown to adopt multiple polymorphic conformations. Recent cryo-electron microscopy studies have identified two predominant fibril polymorphs—designated as the rod and twister forms—which assemble around distinct core regions: the preNAC segment (_47_GVVHGVATVA_56_) and the NACore segment (_68_GAVVTGVTAVA_78_), respectively [30,31]. The structural perturbations at these segments were reported to influence the conformational ensemble of α-Syn fibrils, potentially altering the equilibrium between the fibril polymorphs. Upon glycation, we observed subtle NMR chemical shift changes in the preNAC region, particularly in residues such as H50, G51, and A53. Additional small changes were also observed towards N-terminal region. Consequently, local structure or dynamics of these regions would be perturbed, causing the reduction in α-Syn aggregation propensity upon glycation. Transmission electron microscopy (TEM) provided a direct visual confirmation of these findings. Long, branched amyloid fibrils were observed in DPBS-F, whereas non-fibrillar amorphous assemblies were observed for MGO-F (Figure 1H). Collectively, the biophysical characterization of MGO-F shows that they have a distinct structure from the DPBS-F. However, it is clear that they lose their monomeric structure. Thus, glycation modification of α-Syn causes a change in its aggregation dynamics and characteristics.

To validate if any unbound MGO in MGO-F sample impacted their biophysical characterization, a subset of the MGO-F assemblies was dialyzed to remove any free MGO. Biophysical characteristics of dialysed assemblies remained largely similar to that of non-dialysed MGO-F (Figure S1A to S1C). Further, we did not observe any significant difference (*p*=0.5288) in the N2A cell viability after exposure to MGO-F and dialysed MGO-F with both displaying around 20-25% cell loss after 48 hours of exposure (Figure S1D). This implies that the amount of unbound MGO had a negligible effect on cell viability. This cell loss was very similar to that caused by DPBS-F exposure (22.22%; *p*=0.121 w.r.t. control). Immunostaining for human α-Syn (h-α-Syn) antibody further revelled that in the DPBS-F exposed cells, the assemblies appear to surround the cell, potentially associating with the plasma membrane and only some of it colocalised with cytoskeleton protein β-actin (Figure S1E). After exposure to both dialyzed and non-dialyzed MGO-F, relatively fewer assemblies associated with cell surfaces were visible. This is potentially due to their smaller size, which may have allowed more efficient cellular uptake. Comparable patterns of cell viability loss and cellular accumulation for both dialyzed and non-dialyzed MGO-F also suggest that the presence of unbound MGO did not influence the internalization. Due to challenges associated with the dialysis procedure, such as difficulties in accurately determining fibril concentration, unavoidable protein loss, and issues with maintaining sterility, the further experiments were conducted using non-dialyzed MGO-F.

Collectively, these results suggest that MGO-induced glycation leads to markedly distinct assemblies compared to fibrils, but also show clear modifications relative to the monomers. We, were thus interested to understand the impact of these structurally distinct species in PD pathogenesis *in vivo*.

### 2.2. Glycated α-Syn worsens the behavioral outcomes in mice

Having observed the distinct characteristics of MGO modified and unmodified α-Syn assemblies *in vitro*, we sought to further investigate their effect on PD pathology and progression in mice. To this end, we stereotactically injected randomized male and female mice with either DPBS-F, MGO-F, or DPBS (as control) into the substantia nigra (SN) region of the brain and assessed them behaviourally up to 12 weeks (W) post-surgery (Figure 2A). The open-field test was used to evaluate the general locomotor ability and exploratory behavior of different groups of mice every 4 weeks (Figure 2B). The DPBS-F-injected group displayed a significant reduction in the total distance traveled at the 12W time point (1.5-fold decrease, *p*=0.0005) in comparison to the 4W time point. Similar changes were not observed in other groups.

**Figure 2.**
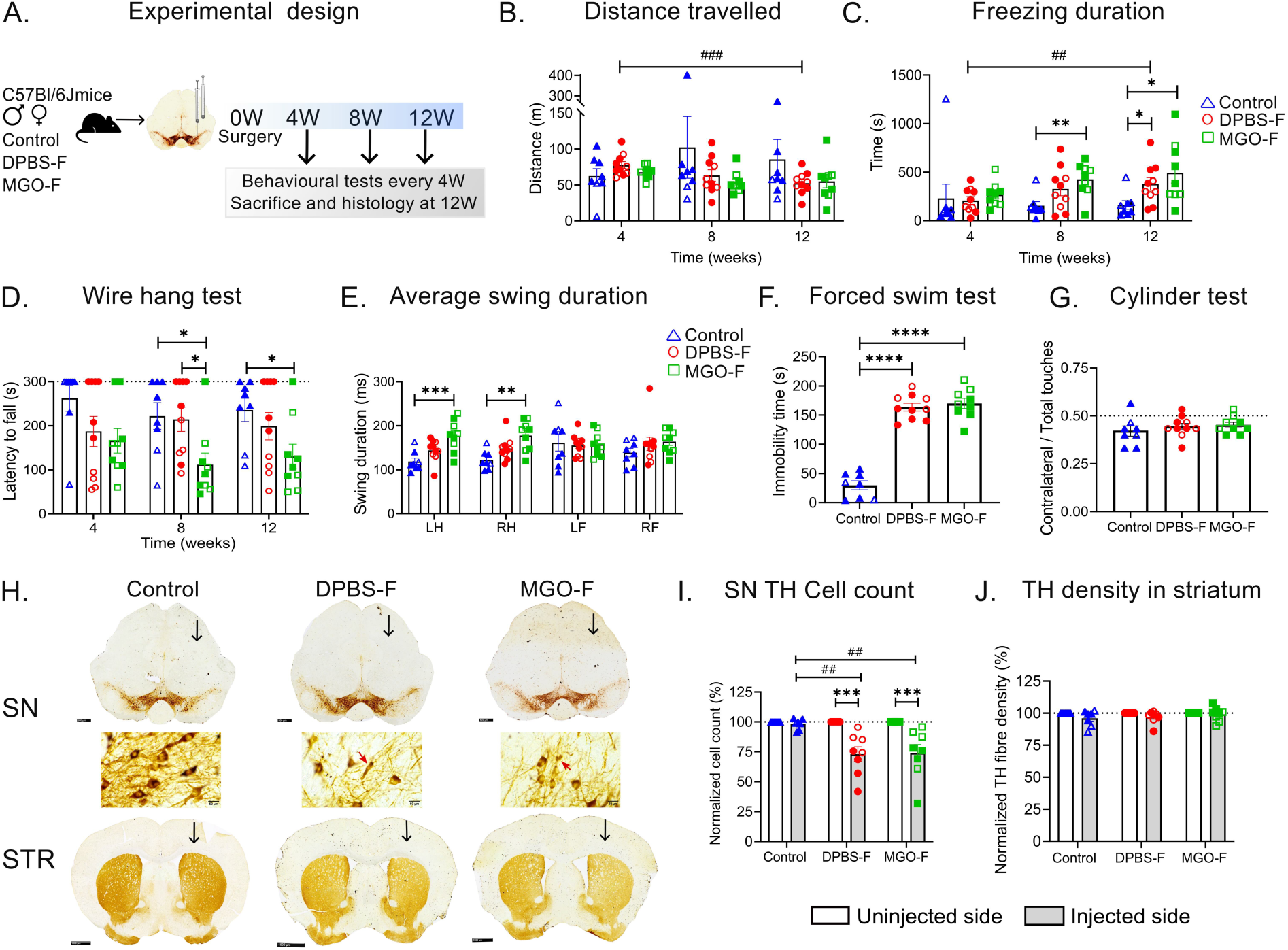
- Glycated α-Syn accelerates behavioral deficits despite similar TH loss in mice. A. Schematic representation of the experimental plan depicting control, DPBS-F, and MGO-F groups sacrificed 12 weeks (12W) post-surgery, with behavioral tests conducted every 4 weeks (4W, 8W and 12W). B. Measure of the total distance traveled in the open field as a test for general locomotor behavior, showing decreased distance traveled by the DPBS-F group with time. C. Increase in freezing duration of the animals in MGO-F sets in at 8W in contrast to 12W in the DPBS-F group. D. Latency to fall in the wire hang test shows strong deterioration in the MGO-F group. E. MGO-F-treated mice showed increased average swing duration in the corridor test, indicative of gait deficits 12 weeks post-surgery (LH-left hindlimb; RH-right hindlimb; LF-left forelimb; RF-right hindlimb). F. Forced swim test for evaluation of depression phenotype at 12W, showing higher immobility in DPBS-F and MGO-F groups. G. Cylinder test to detect preferential paw usage due to unilateral lesion, 12 weeks post-surgery, did not show any motor asymmetry across groups. H. Representative micrographs of SN (scale bar-500 µm) and STR (scale bar-1000 µm) stained for TH at 12W, and SN high magnification (scale bar-20µm), black vertical arrows denote the injected side, red arrows in high magnification images point to the dystrophic neurons in SN. I. Quantification of TH-positive cells in SN shows neurodegeneration in the injected side of DPBS-F and MGO-F. J. Striatal TH fiber density remained unaffected across all three groups. All data are plotted as mean ± SEM, n=8-10 mice in all groups for B to G and n=6-8 mice for I & J. * indicates comparison between groups at a single time point, and ^#^ represents comparison across time within a group. Repeated measures two-way ANOVA for B to D, ordinary two-way ANOVA for E, ordinary one-way ANOVA for F and G, mixed two-way ANOVA for I and J with Tukey’s or Sidak’s multiple comparison test as post-hoc analysis. Filled symbols-female and empty symbols-male in the histograms.

Further, anxiety-like behavior was assessed by measuring the freezing duration in the open field test [32]. DPBS-F injected mice showed a significant increase in freezing duration at the 12W time point in comparison to the control group (2.3-fold increase, *p*=0.038) (Figure 2C). In contrast, MGO-F injected mice displayed a significant increase in the freezing duration in comparison to the control mice as early as 8W (2.8-fold increase, *p*=0.005), which continued to increase further at 12W (3.1-fold increase, *p*=0.036). The DPBS-F-injected group displayed a significant increase in the freezing time at the 12W time point (1.8-fold increase, *p*=0.007) in comparison to the 4W time point. The increase in freezing time was complementary to the decreased distance traveled by DPBS-F mice (Figure 2B). Since control mice did not display any change in freezing duration over time, it rules out the possibility that increasing age or body weight may have impacted the mice’s behavior. Thus, it can be concluded that MGO-F injection caused the earlier onset of non-motor symptoms in the mice.

To assess any changes in the neuromuscular grip strength, a wire hang test was performed. While no significant loss was observed at the 4W timepoint, at 8W, the grip strength of the MGO-F group was decreased compared to the control (*p*=0.039) as well as the DPBS-F group (*p*=0.039) (Figure 2D). At 12W as well, a significant decrease (*p*=0.03) in grip strength was observed in the MGO-F mice compared to the control. In contrast, no such decline was observed in the DPBS-F group in comparison to the control, suggesting exacerbated grip strength loss with MGO-F. Interestingly, male mice in the control group and DPBS-F group performed worse in the wire hang test in comparison to the females (Figure S2A). However, in MGO-F group, both male and female mice show a decreased latency to fall. At 12W, female mice in the MGO-F group had lower latency to fall in comparison to the female mice in the control group (*p*=0.005) and the DPBS-F group (*p*=0.008). In contrast, male mice in any of the groups did not show a significant difference in the latency to fall at 12W. The loss of grip strength is also observed in human PD patients, often as an early symptom occurring years before the onset of classic motor symptoms such as tremors and bradykinesia [33].

Gait abnormalities, another hallmark motor symptom of PD, were also evaluated using the corridor test (Figure 2E). Similar to other tests, MGO-F mice displayed significant gait disturbances 12W post-surgery with increased average swing duration in the left hindlimb (1.5-fold increase; *p*=0.001) and right hindlimb (1.45-fold increase; *p*=0.0018) in comparison to the control group. On the other hand, the DPBS-F group did not show any significant changes in average swing duration in comparison to the control mice. A significant shift in the gait parameters after MGO-F injection indicates that they can cause more severe disruption of motor functions.

In addition to motor symptoms, non-motor symptoms such as depression were assessed using the forced swim test (Figure 2F), as it is a prevalent non-motor symptom faced by PD patients, often before the onset of prominent motor symptoms [34]. As this test can be distressing to the animals, it was conducted only at 12W. Both DPBS-F and MGO-F mice exhibited significantly longer immobility times in comparison to the control mice (5-fold increase; *p*<0.0001), indicating a depressive phenotype. This further supports the anxious phenotype that was observed in the DPBS-F and MGO-F injected mice at 12W. No significant sex-based difference was observed amongst the groups in terms of this trait.

To assess any motor asymmetry induced by a unilateral lesion, the cylinder test was conducted (Figure 2G). The test was conducted only at 12W, as repeated exposure to the cylinder is associated with reduced exploratory behavior and non-performance of the test by the mice. No significant differences in paw preference were observed between the groups, suggesting that the lesion was not severe enough to induce a visible motor asymmetry. This aligns with the fact that significant motor asymmetry in rodents typically requires 70-80% neurodegeneration in one hemisphere [35]. The 12W post-surgery period in the mouse model used here mimics the early to middle stage of PD with a smaller extent of neurodegeneration.

These findings suggest that glycated α-Syn caused a faster onset of motor and non-motor behavioral deficits in comparison to the non-glycated α-Syn. This is in line with the reports that human diabetic PD patients experience more severe motor symptoms with higher cognitive decline compared to non-diabetic PD patients [36,37]. Published literature reports motor, cognitive, and olfactory deficits in MGO-infused Thy-1 α-Syn mice [25]. Further, decreased latency to fall off the rotorod and reduced grip strength are reported in mice injected with D-ribose-glycated α-Syn [27]. While previous studies showed that general glycation in the brain causes these behavioral deficits, we demonstrate that the glycation of α-Syn alone is sufficient in driving more severe behavioral deficits in comparison to non-glycated α-Syn fibrils.

### 2.3. α-Syn assemblies trigger dopaminergic neuronal loss

Mice were sacrificed 12W after the surgery, and their brains were analyzed for various histological markers. TH was used as a marker for dopaminergic neurons and their projections, with a focus on key brain regions relevant to PD - SN and striatum (STR) (Figure 2H). The neuronal loss was ∼25% in the injected side of the DPBS-F (*p*=0.0016) and MGO-F (*p*=0.002) groups in comparison to the injected side of the control mice (Figure 2I). Similarly, both the DPBS-F group (26.8%, *p*=0.0004) and the MGO-F group (26.1%, *p*=0.0006) also showed a similar loss of TH-positive neurons in the injected SN in comparison to their uninjected side. The magnified images of SN displayed several dystrophic dopaminergic neurons in both the α-Syn injected groups. Notably, dopaminergic neuronal loss was observed to be more prominent in female mice compared to males, despite similar behavioral deficits being recorded in most parameters for both MGO-F and DBPS-F groups (Figure S2B). A loss of ∼40% in the TH-positive neurons was noted in the female mice of the DPBS-F and MGO-F groups, in contrast to only ∼15% loss observed in the male mice of the DPBS-F and MGO-F groups. The female mice of the DPBS-F group showed higher TH neuronal cell loss in the SN in comparison to male mice of the same group (*p*=0.035). TH staining at 12W showed no significant change in dopaminergic fiber density between the injected and uninjected sides in either group in the STR, which receives dopaminergic projections from the SN (Figure 2J).

Despite the comparable extent of neurodegeneration in both DPBS-F and MGO-F groups, the more severe behavioral phenotypes observed in the MGO-F group raise questions about the additional factors driving the pathology. Glycated α-Syn may exert its effects through mechanisms beyond direct neurodegeneration, such as neuroinflammation or synaptic dysfunction. In addition, it may be impacting other non-dopaminergic neurons in the basal ganglia circuitry, such as cholinergic, glutamatergic, and GABAergic neurons. Studies have shown that alterations in glutamatergic signaling can precede or accompany dopamine deficiency, potentially contributing to motor deficits and disease severity [25]. Disruptions in the basal ganglia circuitry, even before significant dopamine loss, may underlie the early behavioral changes observed in PD. Further investigations are required to explore the broader impact of glycated α-Syn on different neuronal networks at play.

### 2.4. Both α-Syn assemblies caused accumulation of aggregates in brain

We examined the deposition of α-Syn phosphorylated at serine-129 (p-Syn), a marker for aggregation in these brains. We observed a clear deposition of p-Syn positive aggregates (white arrows) in the SN dopaminergic neurons after injection of both MGO-F and DPBS-F (Figure 3A). Apart from the prominent p-Syn aggregates, thread-like filamentous p-Syn-positive structures reminiscent of Lewy neurites were also evident in a few mice from both groups (yellow arrows). There was a strong increase in p-Syn positive accumulation in the injected side of the DPBS-F group in comparison to the uninjected side (*p*=0.026) as well as with respect to the injected side of control group (p=0.039). However, there was considerable variation in deposition among individual mouse. Around 25% of the mice in the DPBS-F groups displayed a large increase in the deposition compared to the uninjected side (Figure 3B). Remaining mouse brains also had p-Syn deposits, but the increase was small when compared to the uninjected side. Similarly few mouse in the MGO-F group displayed prominent deposition of p-syn but average increase was not statistically significant. In this mild PD model, representing early-to mid-stage PD, excessive p-Syn accumulation is not expected. In addition to the SN, the p-syn deposits were also observed in the STR in both DPBS-F and MGO-F groups (Figure S3). This indicates that despite the glycation modification, α-Syn is still capable of being phosphorylated *in vivo*, and these p-Syn aggregates can propagate to the neuron terminals in STR. This challenges the traditional view that fibrillar structures are the primary drivers of α-Syn toxicity and highlights the need to consider alternative aggregation pathways in PD pathogenesis. Further exploration of these mechanisms is required in the future for a better understanding of the role of glycation in PD progression. The presence of p-Syn in the STR can cause synaptic dysregulations, which may further lead to neuronal dysfunction and promote degeneration [38]. We further investigated if aggregates preferentially deposited in the TH-positive dopaminergic cells. TH-positive neurons in the SN of the DPBS-F group showed a significantly higher colocalization with p-Syn, in comparison to its uninjected side (Mander’s coefficient score 0.28; *p*=0.002) and also in comparison to the injected side of the control (*p*=0.003) (Figure 3C). The preferential accumulation of fibrillar α-Syn aggregates within dopaminergic neurons can lead to their dysfunction, and result in exacerbation of degeneration. In contrast, only some brains in the MGO-F group showed a strong colocalization within the TH-positive neurons of SNc (Mander’s coefficient score 0.18; *p*=0.056). Thus, p-Syn upregulation and its deposition in the dopaminergic neurons of the SN were more prominent in the DPBS-F group.

**Figure 3.**
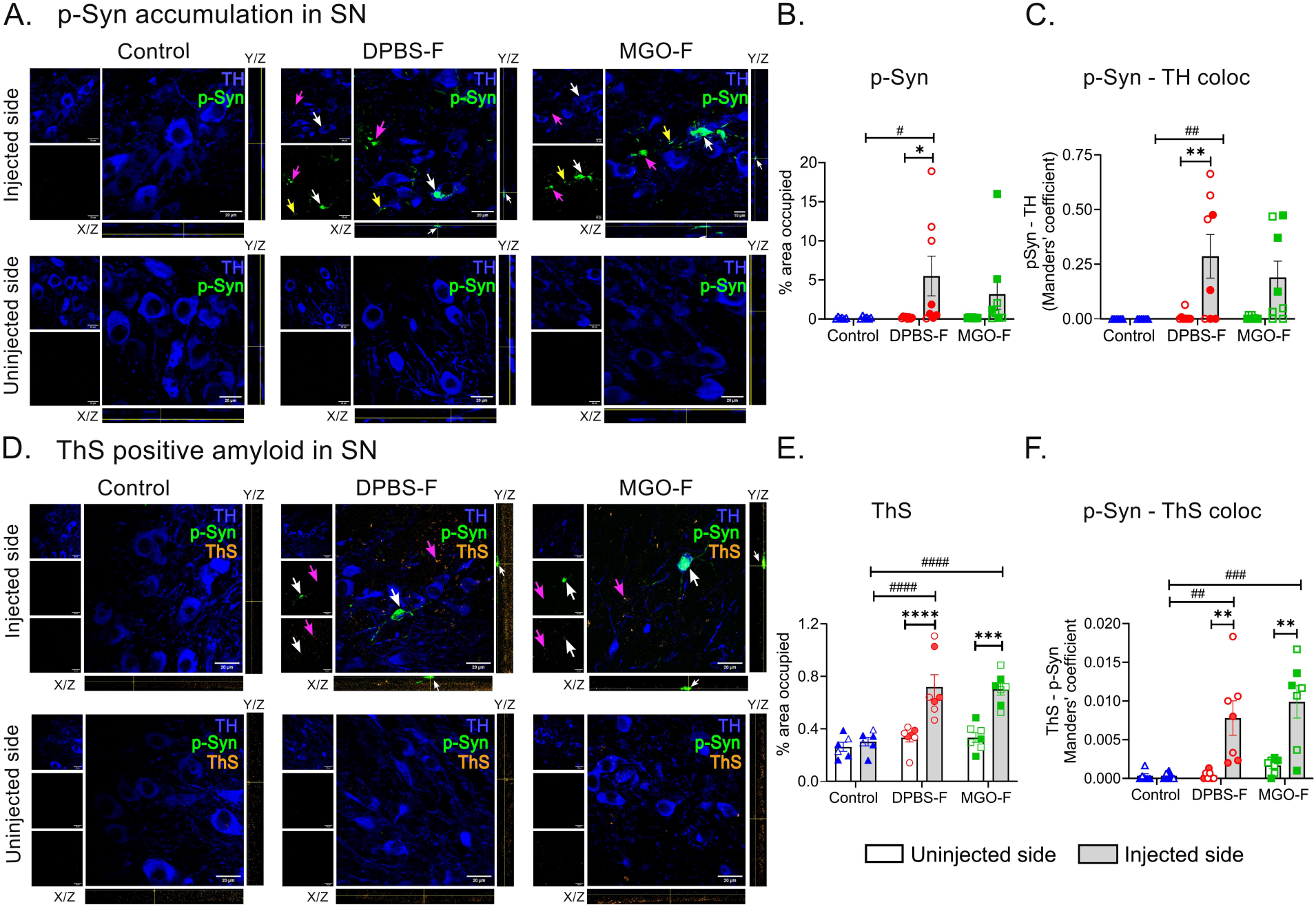
- Accumulation of p-Syn in SN. A. Representative micrographs of SN stained for p-Syn in different groups along with the orthogonal views. White arrows mark the p-Syn aggregates colocalizing with TH, magenta arrows mark p-Syn not localized with TH and yellow arrows mark the Lewy neurites (scale bar-20 µm). B. Quantification of the percentage area occupied by p-Syn in SN. C. Colocalization of p-Syn with TH in SN. D. Representative micrographs of SN showing ThS co-staining with p-Syn and TH. White arrows indicate p-Syn aggregates and ThS deposits in the merged channel with orthogonal views (scale bar-20 µm). E. Quantification of the percentage area occupied by ThS in SN showing increased amyloid deposits in the injected side of DPBS-F and MGO-F, magenta arrows denote non-colocalizing signals. F. Colocalization of p-Syn with ThS in SN showing upregulation in MGO-F and DPBS-F groups. All data are represented mean ± SEM; n= 5-8 mice in each group; mixed two-way ANOVA followed by Tukey’s or Sidak’s multiple comparisons test as post hoc analysis; \**p* < 0.05, **p < 0.01, \*\*\**p* < 0.001, \*\*\*\**p*<0.0001, ^#^*p* < 0.05, ^##^*p* < 0.01, ^###^*p* < 0.001, ^####^*p*<0.0001; * indicates comparison within groups and ^#^ represents comparison between groups; Filled symbols-female and empty symbols-male.

To validate the amyloid nature of the p-syn aggregates, brains were stained for Thioflavin-S (ThS), an amyloid marker (Figure 3D). ThS quantification indicated that the amyloid-positive deposits were abundantly present on the injected side of the DPBS-F (2.2-fold higher, *p*<0.0001) and the MGO-F (2.1-fold higher, *p*=0.0002) groups in comparison to their uninjected sides (Figure 3E). This accumulation was also higher in the injected side of both DPBS-F and MGO-F groups in comparison to the injected side of the control group (*p*<0.0001). A significant increase in colocalization of ThS with p-Syn was observed on the injected side of both the DPBS-F (*p*=0.003) and the MGO-F groups (*p*=0.001) compared to their respective uninjected sides (Figure 3F). This colocalization was significantly higher in the injected side of both DPBS-F (*p*=0.005) and MGO-F (*p*=0.0002) groups in comparison to the injected side of the control group as well. This confirms that amyloid-like deposits can be formed *in vivo* by MGO-F, despite its amorphous nature, further confirming its pathogenic potential. To further characterize the aggregates, their localization with ubiquitin, another protein associated with Lewy bodies and a marker for protein degradation, was examined (Figure S4). Ubiquitin-positive p-Syn aggregates were observed in both DPBS-F and MGO-F groups to a similar extent, indicating that these aggregates are recognized by the cells as pathological and they try to tag them with ubiquitin for degradation attempting to clear them out. However, these ubiquitously labeled, aggregated structures suggest that they evade proteasomal degradation, as is often seen in synucleinopathies [39,40].

These findings demonstrate that glycated α-Syn, though structurally distinct from fibrillar α-Syn, is equally effective in driving Lewy-like pathology. No sex-based differences were exhibited in the accumulation of p-Syn and other hallmark features of these aggregates.

### 2.5. α-Syn assemblies enhance neuroinflammation on the injected side

As dopaminergic neurodegeneration and protein aggregation in both the DPBS-F and MGO-F groups were not markedly different, we investigated additional factors contributing to more severe behavioral deficits observed in the MGO-F group such as the role of microglia and astrocytes, cells known to be involved in maintaining brain homeostasis. Both DPBS-F (1.7-fold increase; *p*=0.002) and MGO-F (1.7-fold increase; *p*=0.013) groups exhibited significantly increased IBA-1 activation as measured by percentage area occupied in the injected side of the substantia nigra compacta (SNc) when compared to their respective uninjected side (Figure 4A, 4B). No significant IBA-1 accumulation was detected in the injected side of the control group compared to its uninjected side, ruling out the role of surgical injury as a cause for this observed neuroinflammation. Significant IBA-1 accumulation was also observed in the injected side of DPBS-F (2.1-fold increase; *p*=0.002) and MGO-F (1.6-fold increase; *p*=0.04) groups in comparison to the injected side of the control group. To assess the immune response mediated by these cells in active clearance of toxic aggregates and debris, colocalization of IBA-1 and p-Syn was examined (Figure 4C). A slight increase, although statistically non-significant, in the colocalization was observed on the injected side of the DPBS-F (Mander’s coefficient score 0.069) and MGO-F (Mander’s coefficient score 0.035) groups compared to their respective uninjected sides (Figure 4D). In addition to the SNc, neuroinflammation in the substantia nigra reticulata (SNr) region was also analyzed (Figure 4E). Statistically significant increase (1.6-fold increase; *p*=0.048) in the levels of IBA-1-positive cells were observed only on the injected side of MGO-F mice in comparison to the injected side of the control group (Figure 4F). These results indicate that MGO-F propagates neuroinflammation more rapidly and extensively compared to DPBS-F, potentially due to its smaller size and altered physicochemical properties. Pronounced inflammation in the SNr, a key region in the basal ganglia motor circuitry, suggests a potential link to the motor deficits observed in the MGO-F group. Microglia, the brain’s resident immune cells, play crucial roles in neuronal homeostasis, repair, and maintenance and are the primary responders to injury or infection. The p-Syn accumulation in the early stages of the disease can put microglia in a chronic inflammatory state. Dysfunctional or persistently activated microglia, in turn, can create an inflammatory microenvironment and enhance neuronal damage.

**Figure 4.**
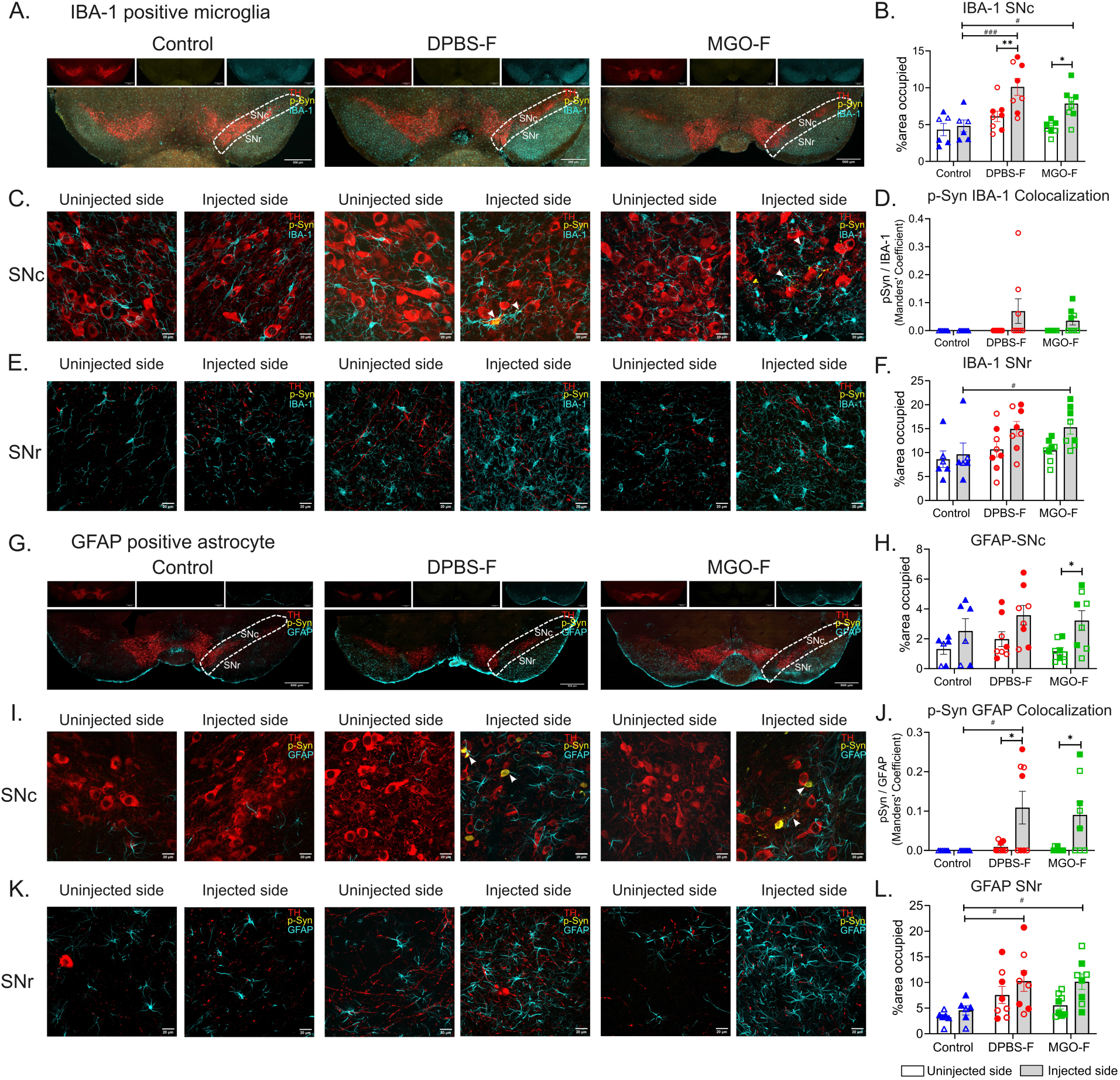
- Both DPBS-F and MGO-F α-Syn assemblies trigger neuroinflammation in the mice brain. A. Representative micrographs of SN co-stained for TH, p-Syn, and IBA-1 (scale bar-500 µm) in indicated groups. B. Quantification of the percentage area occupied by IBA-1 in the SNc region (marked by white dash line in A) shows significant inflammation in the injected side of DPBS-F and MGO-F groups. C. Representative high magnification micrographs of the SNc region depicting co-staining for TH, p-Syn, and IBA-1 (scale bar-20 µm); white arrowheads depict colocalization of IBA-1 with p-Syn. D. Quantification of colocalization of p-Syn with IBA-1 in the SNc region in all three groups. E. Representative higher magnification micrographs of the SNr region (scale bar-20 µm) depicting co-staining for TH, p-Syn, and IBA-1. F. Quantification of the percentage area occupied by IBA-1 in the SNr region, showing enhanced inflammation in the injected side of MGO-F. G. Representative micrographs of SN co-stained for TH, p-Syn and GFAP (scale bar-500 µm) in all three indicated groups. H. Quantification of percentage area occupied by GFAP in the SNc region (marked by white dash line in G) showing increased inflammation in the injected side of MGO-F. I. Representative higher magnification micrographs of the SNc region co-stained for TH, p-Syn and GFAP (scale bar-20 µm); white arrowheads denote colocalization of GFAP with p-Syn. J. Quantification of colocalization of p-Syn with GFAP showing significant upregulation in the injected side of DPBS-F and MGO-F groups. K. Representative higher magnification micrographs of the SNr region co-stained for TH, p-Syn and GFAP (scale bar-20 µm). L. Quantification of the percentage area occupied by GFAP in the SNr region, showing increased inflammation in the injected side of DPBS-F and MGO-F. All data are represented as mean ± SEM; n= 6-8 mice in all groups; Mixed two-way ANOVA followed by Tukey’s or Sidak’s multiple comparisons test as post hoc analysis; \**p* < 0.05, \*\**p* < 0.01, ^#^*p* < 0.05, ^###^*p* < 0.001; * indicates comparison within groups and ^#^ represents comparison between groups; Filled symbols-female and empty symbols-male.

Similarly, activation of GFAP-positive astrocytes was also investigated in the SNc (Figure 4G). A significant increase in the astrocytic activation was observed only in the injected side of MGO-F (2.8-fold increase; *p*=0.032) in comparison to its uninjected side (Figure 4H). A statistically non-significant increase was also detected in the DPBS-F and the control group. Further, colocalization of GFAP with p-Syn was examined and significantly higher scores were noted on the injected side of the DPBS-F (Mander’s coefficient score 0.099; *p*=0.012) and the MGO-F (Mander’s coefficient score 0.087; *p*=0.031) compared to their respective uninjected sides (Figure 4I, 4J). The colocalization scores were higher on the injected side of the DPBS-F group (*p*=0.01) and MGO-F group (*p*=0.037) in comparison to the injected side of the control group as well. This suggests the active involvement of astrocytes in the clearance of p-Syn positive aggregates. While IBA-1 in the SNr region was elevated in only MGO-F groups, GFAP accumulation in the SNr region was more pronounced in the injected side of both DPBS-F (2.3-fold higher; *p*=0.025) and the MGO-F (2.2-fold higher; *p*=0.029) in comparison to the injected side of the control group (Figure 4K, 4L). In line with the more severe TH cell loss, the female mice displayed a slightly higher accumulation of IBA-1 in comparison to the males in both DPBS-F and MGO-F groups. In the SNc region, for the DPBS-F group, a 1.5-fold increase was recorded in males, while a 1.8-fold increase was observed in females (Figure S2C). The injected side of females had significantly higher IBA-1 activation in comparison to the injected side of control female mice (*p*=0.0016) and the control male group (*p*=0.048). In the MGO-F group, a 1.6-fold increase in IBA-1 activation was noted in males, and a 1.8-fold increase was present in females. In the SNr region, the DPBS-F and MGO-F female mice displayed a statistically non-significant mild increase in the IBA-1 accumulation in comparison to the male mice of these groups (data not shown). Although a sex-specific distribution pattern was not observed for the astrocytic activation, the increased accumulation of IBA-1-positive microglia observed in the female mice might promote enhanced SN dopaminergic neurodegeneration compared to the male mice in both the DPBS-F and the MGO-F groups. The exacerbated neuroinflammation can further injure the neurons, paving the way for increased neurodegeneration over time.

To determine whether the inflammation had extended to the distal regions, STR was also examined through IBA-1 and GFAP immunostaining. Interestingly, no significant differences in the microglial activation was observed in any of the groups between the injected and the uninjected sides (Figure S5A, S5B). Similarly, there was no significant accumulation of astrocytic marker, GFAP in the injected side in comparison to the uninjected sides amongst all groups (Figure S5C, S5D). Astrogliosis is associated with chronic neuroinflammation and axonal perturbations, which can exacerbate PD pathophysiology [41]. Activated microglia can further induce astrocytes to release proinflammatory factors, creating a feedback loop that amplifies inflammation and neurodegeneration [42]. The pronounced microglial and astrocytic activation in the SNc and SNr of the MGO-F group might be playing a role in driving the more severe behavioral phenotypes observed in these animals.

### 2.6. MGO-F mediated accumulation of AGE with increased RAGE activation in SN dopaminergic neurons

To understand the mechanisms behind the increased neuroinflammation by glycated α-Syn, the accumulation of AGEs and their receptor, RAGE, was examined in the SN region. AGEs and RAGE are known to play critical roles in inflammation, particularly in metabolic disorders and neurodegenerative diseases [43]. Proteins can be irreversibly modified by AGEs, leading to their dysfunction and disruption of their physiological interactions [44]. AGEs are strongly associated with synucleinopathies in patient brains and are major components of aggregated inclusion bodies [13,22]. RAGE is a multi-ligand receptor (including AGEs, protein aggregates, S100B, HMGB1) that, upon activation, promotes downstream intracellular inflammatory cascades, including NF-kB, MAPK, and ROS [45,46,47]. These pathways can lead to cellular dysfunction and death. Higher levels of these ligands are detected in neurodegenerative diseases [48, 49]. RAGE can be either membrane-bound or present as a soluble receptor in the extracellular matrix or fluid compartments. RAGE is also known to be elevated in the brains of synucleinopathy patients, indicating their potential role in pathology [50].

Despite some animals in both the DPBS-F group and the MGO-F group displayed around a 2-fold increase in AGE levels in comparison to the control mice, the level of AGE in the SNc of the injected side did not show a significant increase in comparison to the uninjected side in any of the treatment groups (Figure 5A, 5B). This may be attributed to the introduction of pre-glycated α-Syn rather than a direct glycating agent such as MGO used in other studies [22,25], which has a broader glycating capacity and could lead to more extensive AGE accumulation. Nevertheless, a significant accumulation of AGEs, specifically in dopaminergic neurons of the DPBS-F and MGO-F groups was observed, indicating vulnerability of dopaminergic neurons to stress with the introduction of MGO-modified or unmodified α-Syn assemblies (Figure 5C). In the MGO-F group (2.3-fold increase; *p*=0.002) as well as the DPBS-F group (2.1-fold increase; *p*=0.016), a prominent colocalization of AGE with TH-positive dopaminergic neurons was observed in the injected side of their SN compared to the injected side of the control group (Figure 5C). Interestingly, the SN neurons in the uninjected sides of both these groups accumulated significantly higher AGE in comparison to those in the uninjected side of the control group. It is possible that the accumulated AGE in the injected side of the brain could have led to increased levels of the same in the uninjected side as well, possibly through interconnected neuronal, vascular, or inflammatory networks.

**Figure 5.**
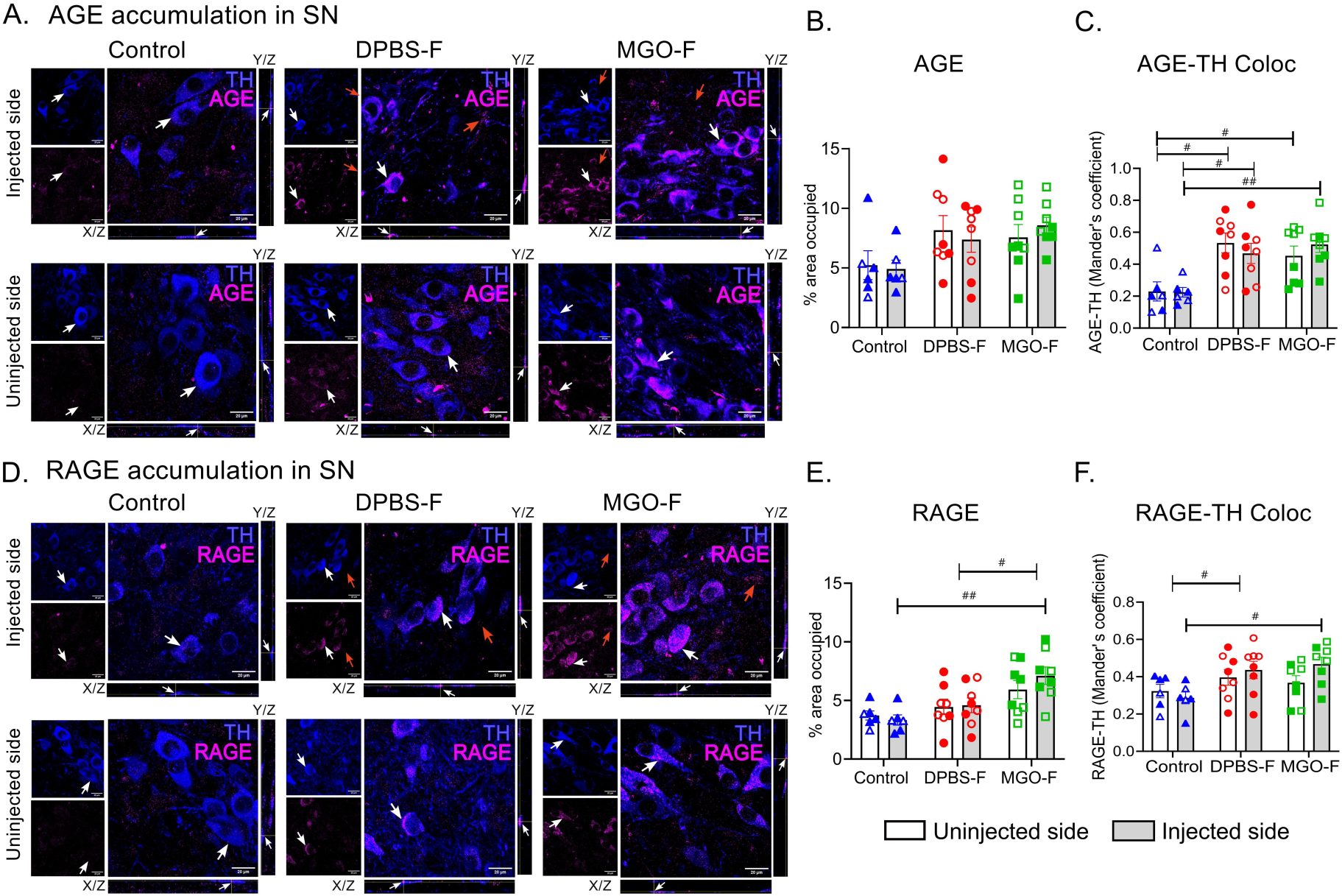
- AGE and RAGE accumulation in SN of mice brain. A. Representative micrographs of SN co-stained for TH and AGE with orthogonal views (scale bar-20 µm). B. Quantification of AGE in the SN depicts no difference between various groups. C. Quantification of TH and AGE colocalization in SN shows upregulation in MGO-F and DPBS-F groups compared to the control. D. Representative micrographs of SN co-stained for TH and RAGE with orthogonal views (scale bar-20 µm). E. RAGE quantification in the SN region, showing enhanced accumulation in the injected side of MGO-F. F. Quantification of TH and RAGE colocalization in SN, confirming increased colocalization with TH in the injected side of MGO-F and DPBS-F groups. White arrows represent colocalizing signals and red arrows denote non-colocalizing signals. All data are represented as mean±SEM; n=6-8 mice in different groups; Mixed two-way ANOVA followed by Tukey’s or Sidak’s multiple comparisons test as post hoc analysis; ^#^*p* < 0.05, ^##^*p* < 0.01; ^#^ represents comparison between groups; Filled symbols-female and empty symbols-male.

Despite the lack of significant AGE upregulation, RAGE expression (Figure 5D, 5E) was markedly elevated in the injected side of the SN of the MGO-F group (2.1-fold increase; *p*=0.001) compared to the injected side of the control group as well as in comparison to the injected side of the DPBS-F group (1.5-fold increase; *p*=0.02). The colocalization of RAGE with TH-positive neurons was significantly higher on the injected side of DPBS-F (1.5-fold increase; *p*=0.039) and MGO-F (1.6-fold increase; *p*=0.010) than on the control’s injected side (Figure 5F). The sex of the animal did not seems to affect the pattern of RAGE accumulation. RAGE, a receptor for both AGE and α-Syn, activates downstream inflammatory pathways upon ligand binding, leading to microglial activation and cytokine release [51]. The aggravated neuroinflammatory response observed in the MGO-F group is likely explained by the enhanced RAGE expression in these animals.

Mechanistically, glycated α-Syn has been shown to enhance the NLRP3 inflammasome pathway, leading to activation of microglia and promotion of neurodegeneration [26]. Further, unmodified α-Syn fibrils activate microglia via binding to the RAGE receptor [52]. It has also been demonstrated that better survival of dopaminergic neurons was observed in siRNA-induced RAGE-deficient mice compared to the wild-type control in an MPTP mouse model [53]. Moreover, targeted inhibition of RAGE reverses α-Syn accumulation and neuroinflammation [54], suggesting that targeting RAGE signaling could be considered a promising therapeutic strategy for mitigating glycation-induced PD pathology.

## 3. CONCLUSIONS

Our study provides a detailed comparison of how glycation modifies α-Syn into distinct assemblies, leading to differential behavioral outcomes, inflammatory responses, and aggregation patterns in the mouse brain. While multiple factors involved in diabetes contribute to the increased risk of neurodegeneration, we have highlighted the contribution of glycated α-Syn in this study. We demonstrate that glycated α-Syn is sufficient to exacerbate PD behavioral symptoms despite causing similar dopaminergic neurodegeneration in comparison to non-glycated fibrils. Such a comparison of motor progression to examine the PD symptoms resulting from MGO-F injection in comparison to the unmodified fibrils has not been described in the literature before. While our study explored the neurodegeneration at early stages, it is possible that over a longer duration, exacerbated neurodegeneration may be caused by glycated α-Syn.

The AGE modifications induced by MGO on α-Syn promote neuroinflammation in the MGO-F group via RAGE activation to a larger extent than the α-Syn fibrils. Besides the SNc, the SNr, which plays major role in regulating the basal ganglia circuitry, was also observed to have higher neuroinflammation after MGO-F exposure. The effects of glycated α-Syn in other brain regions have not been closely examined in previous studies. Higher microglial activation and, consequently, more pronounced dopaminergic neurodegeneration were exhibited by female mice compared to male mice. Thus, with exacerbated glycation of α-Syn, there is also a potential risk for early-onset of PD symptoms and pathology, particularly in diabetes patients. This is concerning given that the average PD onset age for the Indian PD population is already a decade younger than in the Western populations [55]. The increased toxicity of glycated α-syn assemblies underscores their critical role in PD pathogenesis and positions them as a potential candidate for therapeutic targeting.

## 4. MATERIALS AND METHODS

### 4.1. Protein expression and purification

The pT7-7 plasmid encoding wild-type human α-Syn (kind gift from Dr. Nixon Abraham, IISER Pune) was used to transform *Escherichia coli* BL21 (DE3) competent cells. After induction by 1 mM isopropyl β-d-1-thiogalactopyranoside (IPTG) for 4 hrs, cells were centrifuged and the cell pellet was suspended in a buffer containing 50 mM Tris-Cl pH 7.4, 500 mM NaCl, 10 mM EDTA with protease inhibitor cocktail. Further, cell suspension was heated at 99°C for 20 minutes, followed by pulse sonication and subsequently centrifuged to remove cell debris. The resulting supernatant was subjected to ammonium sulphate fractionation at 4°C. Resulting protein pellet was dissolved in desalting buffer (0.5MM Tris-Cl pH 7.4, 2M NaCl, 0.5M EDTA), and loaded onto a HiPrep Desalting 20 mL column connected to an Akta Pure FPLC system pre-equilibrated with desalting buffer. The protein was eluted at a flow rate of 1 mL/min and then loaded onto a HiPrep Q FF column equilibrated with an elution buffer (10 mM Tris-Cl pH 7.4, 500 mM NaCl). Fractions containing α-Syn were pooled and concentrated using a Vivaspin 3 kDa concentrator. Protein concentration was determined using a NanoDrop 1000 spectrophotometer (Thermo Scientific) at 280 nm with a theoretical extinction coefficient of 5960 M⁻¹ cm⁻¹[56].

### 4.2. Aggregation/fibrillation of α-synuclein

α-Syn (5 mg/mL in DPBS) solution was first centrifuged to remove any potential aggregates and then incubated at 37°C for 96 hours with continuous shaking at 950 rpm to produce fibrils (DPBS-F). In parallel, the reaction mixture was also incubated with 5 mM purified MGO (Sigma Aldrich) under similar conditions. Fibril formation was monitored every 24 hours by measuring ThT fluorescence using a TECAN Infinite M200 Pro spectrofluorometer. A triplicate of the reaction mixture was prepared in dark 96-well plates (Corning Incorporated), with each well containing 3.5 µM protein sample and 10 µM ThT (Sigma Aldrich) diluted in DPBS. Mean fluorescence readings were recorded at 37°C with excitation and emission wavelength set at 440 nm and 480 nm respectively. α-Syn aggregates formed in presence of MGO (MGO-F) was dialyzed to remove any free MGO from the mixture using a dialyzing membrane unit (Thermo Fisher Scientific).

### 4.3. Far-UV Circular Dichroism (CD) Spectroscopy

Far-UV CD spectra of 20 µM aggregated α-Syn (DPBS-F and MGO-F) and monomeric control were recorded in DPBS using a BioLogic MOS-500 CD Spectrometer at 20°C. Each spectrum represents the average of three scans.

### 4.4. Dynamic Light Scattering (DLS)

Hydrodynamic radii of 70 µM α-Syn aggregates (DPBS-F and MGO-F) and monomeric control were analysed by DLS using a Wyatt Technology DynaPro NanoStar instrument at 25°C, using the DYNAMICS software. For each sample, the correlation of percent total mass in solution versus particle radii was determined as the average of 10 acquisitions.

### 4.5. Transmission Electron Microscopy (TEM)

A 10 µM solution of aggregated α-Syn (DPBS-F and MGO-F) and monomeric control was applied to carbon-coated grids followed by 3% uranyl acetate staining and subsequent washes with deionized sterile water. The samples were imaged using a FEI TECNAI G2 Spirit BioTwin 120 kV TEM microscope.

### 4.6. Matrix-Assisted Laser Desorption/Ionization Time-of-Flight (MALDI-TOF) Mass Spectrometry

50 µM monomeric control and α-Syn aggregates (DPBS-F and MGO-F) were prepared in DPBS. The SDHB matrix (Bruker Daltonics) and each sample were mixed in a 1:1 ratio using TA30 buffer (3:7 acetonitrile and 0.1% trifluoroacetic acid). The mixture was spotted onto steel target plates and analyzed using a Bruker Ultraflextreme MALDI-TOF/TOF mass spectrometer.

### 4.7. Heteronuclear Single Quantum Coherence Nuclear Magnetic Resonance (HSQC-NMR)

NMR measurements were conducted at 5°C using Bruker Avance-III spectrometers operating at a ¹H frequency of 700 MHz, equipped with a triple resonance (TXI) probe with a z-axis gradient. NMR data were processed using NMRPipe, and peak assignments were performed using Sparky software. NMR samples of ¹⁵N-labeled α-Syn were prepared at a concentration of 350 µM in 20 mM phosphate buffer (pH 7.4) with 0.05% sodium azide (NaN) and 5% deuterium oxide (D_2_O). To monitor the effect of MGO on α-Syn aggregation, ^1^H-^15^N HSQC spectra of α-Syn were recorded 96 hours post-incubation with and without MGO under fibrillation conditions and compared with monomeric control. Absolute intensity values were plotted and overlaid to assess the degree of intensity differences amongst the monomer, DPBS-F, and MGO-F using OriginPro.

### 4.8. Cell culture and immunocytochemistry

N2a cells were cultured in DMEM media (Invitrogen) supplemented with 10% fetal bovine serum (Gibco). Cells were maintained at 37°C in a humidified incubator with a 5% CO_2_ atmosphere. 4×10³ cells/well were seeded in triplicate onto 96-well plates. Different α-Syn species were diluted to a final concentration of 10 µM in DPBS, bath sonicated (30 seconds ON, 30 seconds OFF, total 60 minutes), and added to the cell culture medium 24 hours after seeding. Cell viability was tested using the MTT (3-(4,5-Dimethylthiazol-2-yl)-2,5-Diphenyltetrazolium Bromide) assay. After incubating the cells for 3 hours at 37°C with MTT solution, methanol was used to solubilize the MTT crystals, and absorbance at 570 nm was measured using a TECAN Infinite M200 Pro plate reader. The average cell viability was calculated relative to the untreated control.

For immunofluorescence studies, N2a cells were cultured onto 13-mm coverslips in 24-well plates and treated as described earlier. 48 hours post-incubation, they were fixed with 4% paraformaldehyde (PFA). After washes with PBS, blocking was carried out with PBS-TB buffer (DPBS containing 0.01% Triton X-100 and 1% BSA) at room temperature (RT). This was followed by incubation with primary antibodies for 3 hours at RT. After washing, the fluorophore-conjugated secondary antibodies were added and incubated for 2 hours. Cells were then stained with 0.3 µM DAPI for 1 minute and mounted using ProLong™ Gold Antifade Mountant (Invitrogen). Images were acquired using an Olympus inverted confocal laser scanning microscope (FLUOVIEW FV3000).

### 4.9. Stereotaxic surgery in mice

All procedures were approved by the Institutional Animal Ethics Committee (IAEC) according to the guidelines laid by the Committee for the Purpose of Control and Supervision of Experiments on Animals (CPCSEA), India. Around 12 weeks old male and female C57Bl/6J mice were randomly assigned to the following experimental groups: control, DPBS-F, and MGO-F. Stereotaxic surgeries were performed in the above-mentioned groups using a protocol described earlier [57]. Briefly, mice received 1 µL of either freshly sonicated (1s ON-1s OFF, total for 1 min) MGO-F (5 mg/mL) or DPBS-F (5 mg/mL) at a flow rate of 250 nL/min. The control group received an equivalent volume of 1X DPBS. The materials were injected into the medial SN (−1.1 mm ML, −4.15 mm DV, −3.2 mm AP) and the lateral SN (−1.7 mm ML, −4.6 mm DV, −3.2 mm AP). The coordinates were labeled relative to the bregma. The syringe was held in place for an additional 5 minutes post-injection to prevent backflow. All the mice were housed in the animal facility at IISER-Thiruvananthapuram, under a 12-hour light/dark cycle, with ad-libitum access to chow feed and water.

### 4.10. Behavioral analysis

Behavioral tests were conducted every 4 week post-surgery upto 12 weeks. Animals were acclimatized to the sound-attenuated behavior room for at least 30 minutes before testing.

#### Open field test

Animals were placed in a square arena (50 cm × 50 cm × 50 cm) with opaque walls and allowed to explore for 30 minutes (5 minutes of acclimatization followed by 25 minutes of testing). Their movements were video recorded, and various locomotor parameters were extracted using ANY-maze software.

#### Corridor test

Mice were allowed to walk through a transparent glass corridor (100 cm in length), and their movement was recorded using a camera placed at the bottom. Gait parameters were extracted using Visual Gait Lab software [58].

#### Wire hang test

The mice were suspended from the top of a wire cage at a height of 40 cm, with a soft bedding cushion placed below to prevent injury upon a fall. The latency to fall was recorded, with a maximum cutoff time of 300 seconds.

#### Forced swim test

Mice were gently introduced into a water-filled cylinder. They were video recorded for 6 minutes (2 minutes of acclimatization followed by 4 minutes of analysis). The time spent without any attempt to swim or move was taken as the total immobility time.

#### Cylinder test

Mice were placed in a transparent glass cylinder, and their exploratory behavior was recorded for up to 5 minutes. The first 30 paw touches were analyzed to calculate the ratio of contralateral versus ipsilateral paw touches on the cylinder wall. Paw touches to the bottom of the cylinder were excluded.

### 4.11. Histology

Mice were euthanized 12 week post-surgery by cervical dislocation and transcardially perfused with ice-cold 1X DPBS followed by 4% PFA. Brains were then post-fixed in 4% PFA for 24 hours, cryoprotected in 20% sucrose, and sectioned coronally at 30 µm thickness using a vibratome (Leica VT1000 S). The STR and SN sections were stored in DPBS with 0.02% sodium azide at 4°C. Free-floating sections were permeabilized with 0.1% Triton X-100 for 30 minutes at RT. Antigen retrieval was performed in citrate buffer (pH 6.0) at 80°C for 30 minutes, followed by blocking with 2% BSA for 1 hour at RT. The sections were incubated with primary antibodies overnight at 4°C. After washes, secondary antibody (HRP-or AlexaFluor-conjugated) incubation for 6 hours at RT or 4°C was done. For fluorescent staining, sections were washed and mounted with Vectashield Antifade Mounting Medium (H-1900). For ThS staining, after fluorescent staining, the slices were incubated with 0.01% ThS solution (Sigma) and incubated at RT for 10 minutes, followed by multiple ethanol rinses and DPBS washes [59]. For chromogenic staining, the sections were further incubated with ABC reagent (Vectastain ABC Kit) and developed with 0.05% DAB (Sigma Aldrich) in 0.01% H_2_O_2_ (HiMedia) and mounted using DPX mountant (HiMedia) [60]. Antibody details are provided in Table 1.

**Table-1.**
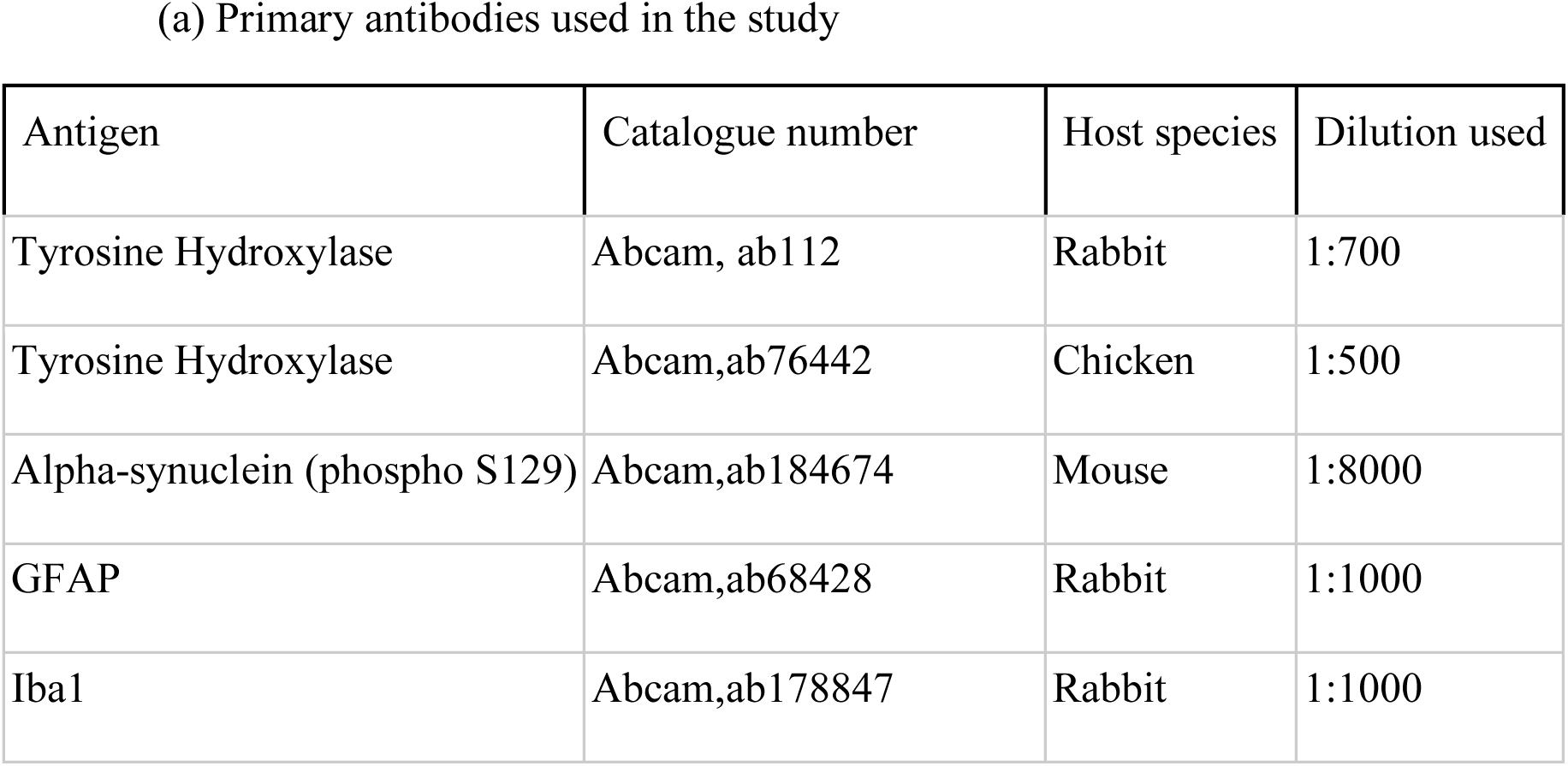
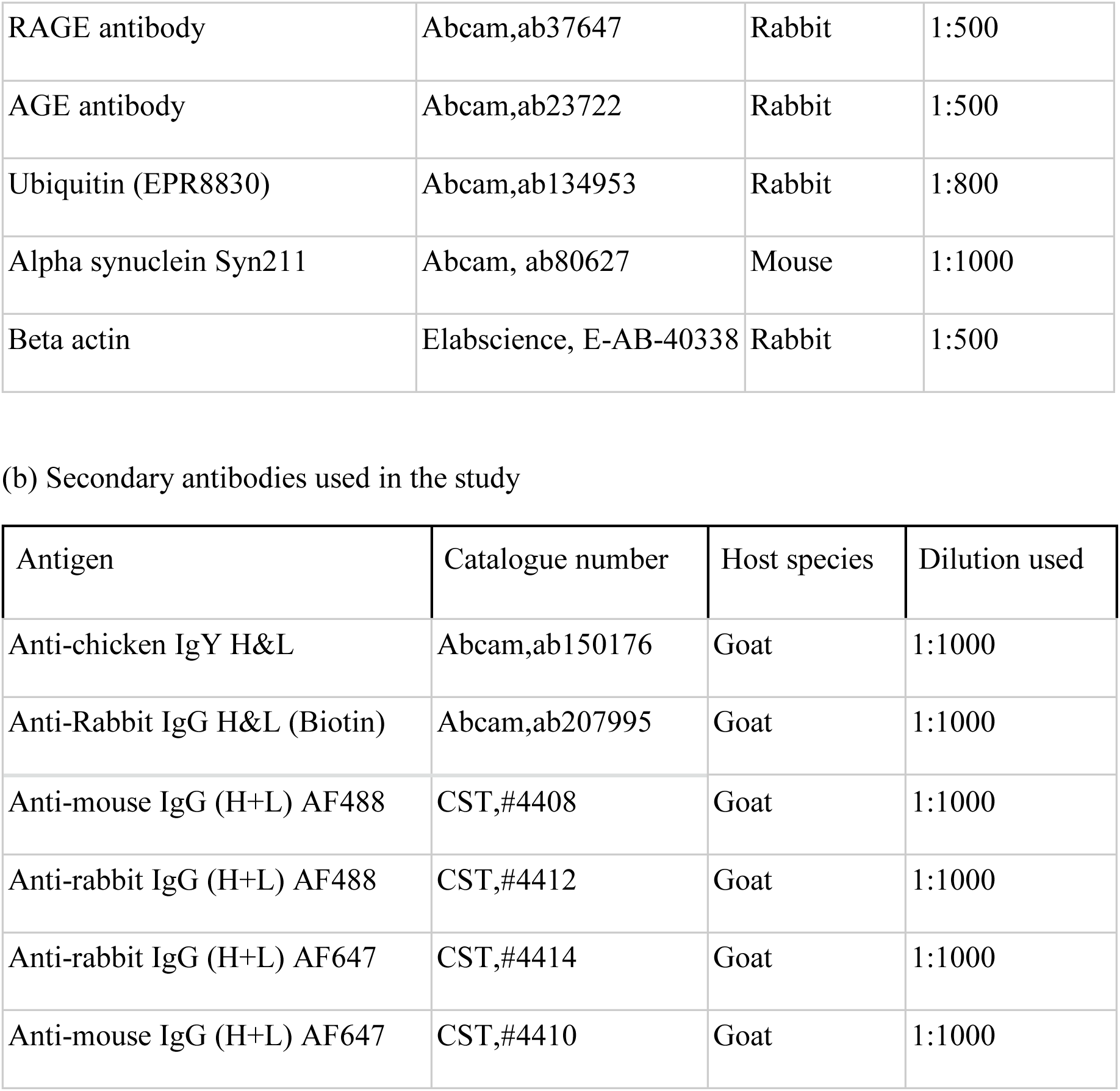

| Antigen | Catalogue number | Host species | Dilution used |
| --- | --- | --- | --- |
| Tyrosine Hydroxylase | Abcam, ab112 | Rabbit | 1:700 |
| Tyrosine Hydroxylase | Abcam, ab76442 | Chicken | 1:500 |
| Alpha-synuclein (phospho S129) | Abcam, ab184674 | Mouse | 1:8000 |
| GFAP | Abcam, ab68428 | Rabbit | 1:1000 |
| Iba1 | Abcam, ab178847 | Rabbit | 1:1000 |
| RAGE antibody | Abcam,ab37647 | Rabbit | 1:500 |
| AGE antibody | Abcam,ab23722 | Rabbit | 1:500 |
| Ubiquitin (EPR8830) | Abcam,ab134953 | Rabbit | 1:800 |
| Alpha synuclein Syn211 | Abcam, ab80627 | Mouse | 1:1000 |
| Beta actin | Elabscience, E-AB-40338 | Rabbit | 1:500 |

(b) Secondary antibodies used in the study
| Antigen | Catalogue number | Host species | Dilution used |
| --- | --- | --- | --- |
| Anti-chicken IgY H&L | Abcam,ab150176 | Goat | 1:1000 |
| Anti-Rabbit IgG H&L (Biotin) | Abcam,ab207995 | Goat | 1:1000 |
| Anti-mouse IgG (H+L) AF488 | CST,#4408 | Goat | 1:1000 |
| Anti-rabbit IgG (H+L) AF488 | CST,#4412 | Goat | 1:1000 |
| Anti-rabbit IgG (H+L) AF647 | CST,#4414 | Goat | 1:1000 |
| Anti-mouse IgG (H+L) AF647 | CST,#4410 | Goat | 1:1000 |

### 4.12. Imaging and Quantification

DAB-stained SN and STR were imaged at 10X and 4X magnification respectively using bright-field microscopy. Fluorescent-stained slices were imaged at 20X magnification using Olympus upright fluorescence microscope (BX63) or Leica upright fluorescence microscope, while 60X and 100X images were acquired using a Fluoview FV3000 inverted confocal microscope. Dopaminergic neuronal cell bodies in the SN were analyzed using the cell counter plugin in ImageJ after drawing a region of interest outlining the SN region. For each mouse, 6-8 brain sections spread across the rostro-caudal extent of SN were used for counting. The dopaminergic fiber density in the STR was assessed by measuring mean intensity using ImageJ. For each mouse, 8-10 brain sections spread across the rostro-caudal extent of STR were used for quantification. For IBA-1 and GFAP analysis, 20X (in case of SNc and SNr) and 10X (in case of STR) images were used to delineate the region of interest, and either the percentage area occupied by the signal as shown previously [61] or Mander’s coefficient (for colocalization quantifications) was calculated using ImageJ JaCoP plugin as shown previously [62]. For quantification of other signals, three 100X images were captured along the medio-lateral extent of SN per hemisphere from one slice of each animal’s brain across all groups for the analysis.

### 4.13. Statistical analysis

All data were plotted and analyzed using GraphPad Prism 9.3.1. Data points are represented as mean ± standard error of the mean (SEM). Statistical comparisons were performed using either one-way ANOVA followed by Tukey’s multiple comparison post-hoc analysis or two-way mixed-design ANOVA to examine the effects between treatment groups and within groups (left and right hemispheres). If interactions after the simple effects were significant, they were followed by Tukey’s multiple comparison post-hoc analysis. When they were non-significant, main effects were interpreted independently, followed by Tukey’s multiple comparison post-hoc analysis for between-group effects and Sidak’s multiple test for within-group effects.

## Author Information

Akshaya Rajan - School of Biology, IISER-Thiruvananthapuram, Kerala, India-695551

Shaliya P H - School of Biology, IISER-Thiruvananthapuram, Kerala, India-695551

Anish Varghese - School of Biology, IISER-Thiruvananthapuram, Kerala, India-695551

Ann Teres Babu - School of Chemistry, IISER-Thiruvananthapuram, Kerala, India-695551

Vinesh Vijayan - School of Chemistry, IISER-Thiruvananthapuram, Kerala, India-695551

## Author Contributions

P.T. planned and supervised the research, obtained funding, and co-wrote the manuscript. A.R. conducted the research, analyzed data, and co-wrote the manuscript. A.V. standardized and performed glycation and biophysical characterization experiments. S.P.H. contributed to the cell culture experiments. A.T.B. and V.V. helped with NMR experiments and data analysis. All authors read and approved the final draft of the manuscript.

## Funding Sources

P.T. is funded by grants from Science and Engineering Research Board, India (SRG/2021/000981), DBT/Wellcome Trust India Alliance Early Career Fellowship (IA/E/17/1/503664), Cure Parkinson’s Trust, UK (PT01), and intramural funds from IISER-Thiruvananthapuram. We also acknowledge the FIST funding for Animalium, IISER-Thiruvananthapuram. A.R. acknowledges fellowship support from the Department of Biotechnology, Government of India.

## Acknowledgment

We thank Santhosh Kumar and Unnati Agrawal for helping with the animal experiments. We also thank Shreshth Shekhar for help with the gait analysis. We thank Dr. Nixon M. Abraham (IISER Pune), Dr. Murthy Srinivasula (IISER-Thiruvananthapuram) and Dr. Vijay Jayaraman (IISER-Thiruvananthapuram) for sharing the resources. We also extend our thanks to Dr. Nongmaithem Sadananda Singh (IISER-Thiruvananthapuram) for sharing the cell culture facility.

## ABBREVIATIONS

PD: Parkinson’s disease
STR: Striatum
SNc: Substantia nigra compacta
SNr: Substantia nigra reticulata
MGO: Methylglyoxal
AGE: Advanced glycation end products
RAGE: Receptor for advanced glycation end products
ThT: Thioflavin-T
DLS: Dynamic light scattering
CD: Circular dichroism
MALDI-TOF/MS: Matrix-assisted laser desorption ionization-time of flight mass spectrometry HSQC-NMR Heteronuclear Single Quantum Coherence-Nuclear magnetic resonance MTT 3-(4,5-dimethylthiazol-2-yl)-2,5-diphenyltetrazolium bromide
N2a: Neuro-2a
TH: Tyrosine hydroxylase
IBA1: Ionized calcium binding adaptor molecule 1
GFAP: Glial fibrillary acidic protein
p-Syn: Phosphorylated alpha synuclein Serine-129
Ubq: Ubiquitin
ThS: Thioflavin-S

## Notes

### Competing Interest Statement

The authors have declared no competing interest.

## REFERENCES

[1] W. Poewe et al., “Parkinson disease,” Nature Reviews Disease Primers, vol. 3, pp. 1–21, 2017, doi: 10.1038/nrdp.2017.13.

[2] E. R. Dorsey et al., “Global, regional, and national burden of Parkinson’s disease, 1990– 2016: a systematic analysis for the Global Burden of Disease Study 2016,” The Lancet Neurology, vol. 17, no. 11, pp. 939–953, Nov. 2018, doi: 10.1016/S1474-4422(18)30295-3.

[3] D. Emin et al., “Small soluble α-synuclein aggregates are the toxic species in Parkinson’s disease,” Nat Commun, vol. 13, no. 1, p. 5512, Sep. 2022, doi: 10.1038/s41467-022-33252-6.

[4] P. Bhattacharjee, A. Öhrfelt, T. Lashley, K. Blennow, A. Brinkmalm, and H. Zetterberg, “Mass Spectrometric Analysis of Lewy Body-Enriched α-Synuclein in Parkinson’s Disease,” J. Proteome Res., vol. 18, no. 5, pp. 2109–2120, May 2019, doi: 10.1021/acs.jproteome.8b00982.

[5] V. A. Petyuk et al., “Proteomic Profiling of the Substantia Nigra to Identify Determinants of Lewy Body Pathology and Dopaminergic Neuronal Loss,” J. Proteome Res., vol. 20, no. 5, pp. 2266–2282, May 2021, doi: 10.1021/acs.jproteome.0c00747.

[6] I. Martinez-Valbuena et al., “α-Synuclein molecular behavior and nigral proteomic profiling distinguish subtypes of Lewy body disorders,” Acta Neuropathol, vol. 144, no. 2, pp. 167–185, Aug. 2022, doi: 10.1007/s00401-022-02453-0.

[7] L. Patterson, S. P. Rushton, J. Attems, A. J. Thomas, and C. M. Morris, “Degeneration of dopaminergic circuitry influences depressive symptoms in Lewy body disorders,” Brain Pathology, vol. 29, no. 4, pp. 544–557, 2019, doi: 10.1111/bpa.12697.

[8] N. Zhao et al., “Quality of life in Parkinson’s disease: A systematic review and meta-analysis of comparative studies,” CNS Neuroscience & Therapeutics, vol. 27, no. 3, pp. 270–279, 2021, doi: 10.1111/cns.13549.

[9] M. C. Hardenberg et al., “Observation of an α-synuclein liquid droplet state and its maturation into Lewy body-like assemblies,” Journal of Molecular Cell Biology, vol. 13, no. 4, pp. 282–294, Apr. 2021, doi: 10.1093/jmcb/mjaa075.

[10] N. Candelise et al., “Effect of the micro-environment on α-synuclein conversion and implication in seeded conversion assays,” Transl Neurodegener, vol. 9, no. 1, p. 5, Dec. 2020, doi: 10.1186/s40035-019-0181-9.

[11] A. Farzadfard et al., “Glycation modulates alpha-synuclein fibrillization kinetics: A sweet spot for inhibition,” Journal of Biological Chemistry, vol. 298, no. 5, May 2022, doi: 10.1016/j.jbc.2022.101848.

[12] E. Vasili, et al., “Glycation of alpha-synuclein enhances aggregation and neuroinflammatory responses,” Jul. 02, 2024, bioRxiv. doi: 10.1101/2024.06.27.600956.

[13] G. Münch et al., “Crosslinking of α-synuclein by advanced glycation endproducts — an early pathophysiological step in Lewy body formation?,” Journal of Chemical Neuroanatomy, vol. 20, no. 3–4, pp. 253–257, Dec. 2000, doi: 10.1016/S0891-0618(00)00096-X.

[14] R. Liu et al., “Fluorescent advanced glycation end products in type 2 diabetes and its association with diabetes duration, hemoglobin A1c, and diabetic complications,” Front. Nutr., vol. 9, Dec. 2022, doi: 10.3389/fnut.2022.1083872.

[15] Y. W. Yang et al., “Increased risk of Parkinson disease with diabetes mellitus in a population-based study,” Medicine, vol. 96, no. 3, 2017, doi: 10.1097/MD.0000000000005921.

[16] J. L. Y. Cheong, E. de Pablo-Fernandez, T. Foltynie, and A. J. Noyce, “The Association Between Type 2 Diabetes Mellitus and Parkinson’s Disease,” Journal of Parkinson’s Disease, vol. 10, no. 3, pp. 775–789, 2020, doi: 10.3233/jpd-191900.

[17] A. Konig, H. V. Miranda, and T. F. Outeiro, “Alpha-Synuclein Glycation and the Action of Anti-Diabetic Agents in Parkinson’s Disease,” Journal of Parkinson’s Disease, vol. 8, no. 1, pp. 33–43, Jan. 2018, doi: 10.3233/JPD-171285.

[18] T. Li, H. Cao, and D. Ke, “Type 2 Diabetes Mellitus Easily Develops into Alzheimer’s Disease via Hyperglycemia and Insulin Resistance,” CURR MED SCI, vol. 41, no. 6, pp. 1165–1171, Dec. 2021, doi: 10.1007/s11596-021-2467-2.

[19] Y. Tang, P. Zhang, L. Li, and J. Li, “Diabetic Peripheral Neuropathy and Glycemia Risk Index in Type 2 Diabetes: A Cross-Sectional Study,” DMSO, vol. Volume 17, pp. 4191–4198, Nov. 2024, doi: 10.2147/DMSO.S482824.

[20] K. Barinova, M. Serebryakova, E. Sheval, E. Schmalhausen, and V. Muronetz, “Modification by glyceraldehyde-3-phosphate prevents amyloid transformation of alpha-synuclein,” Biochimica et Biophysica Acta (BBA) - Proteins and Proteomics, vol. 1867, no. 4, pp. 396–404, Apr. 2019, doi: 10.1016/j.bbapap.2019.01.003.

[21] A. B. Uceda, J. Frau, B. Vilanova, and M. Adrover, “Glycation of α-synuclein hampers its binding to synaptic-like vesicles and its driving effect on their fusion,” Cellular and Molecular Life Sciences 2022 79:6, vol. 79, no. 6, pp. 1–19, Jun. 2022, doi: 10.1007/S00018-022-04373-4.

[22] H. Vicente Miranda et al., “Glycation potentiates α-synuclein-associated neurodegeneration in synucleinopathies,” Brain, vol. 140, no. 5, pp. 1399–1419, May 2017, doi: 10.1093/BRAIN/AWX056.

[23] L. Mariño, A. Belén Uceda, F. Leal, and M. Adrover, “Insight into the Effect of Methylglyoxal on the Conformation, Function, and Aggregation Propensity of α-Synuclein,” Chemistry – A European Journal, vol. 30, no. 36, p. e202400890, 2024, doi: 10.1002/chem.202400890.

[24] Y. Q. Lv et al., “Long-term hyperglycemia aggravates α-synuclein aggregation and dopaminergic neuronal loss in a Parkinson’s disease mouse model,” Translational Neurodegeneration, vol. 11, no. 1, pp. 1–16, Dec. 2022, doi: 10.1186/S40035-022-00288-Z/FIGURES/7.

[25] A. Chegão et al., “Glycation modulates glutamatergic signaling and exacerbates Parkinson’s disease-like phenotypes,” npj Parkinson’s Disease 2022 8:1, vol. 8, no. 1, pp. 1–22, Apr. 2022, doi: 10.1038/s41531-022-00314-x.

[26] M. Kumari, K. S. Bisht, K. Ahuja, R. K. Motiani, and T. K. Maiti, “Glycation Produces Topologically Different α-Synuclein Oligomeric Strains and Modulates Microglia Response via the NLRP3-Inflammasome Pathway,” ACS Chem. Neurosci., vol. 15, no. 20, pp. 3640–3654, Oct. 2024, doi: 10.1021/acschemneuro.4c00057.

[27] M. Kumari, B. Kumar, K. S. Bisht, and T. K. Maiti, “Glycation renders ɑ-synuclein oligomeric strain and modulates microglia activation,” bioRxiv, p. 2022.01.15.476311, Jan. 2022, doi: 10.1101/2022.01.15.476311.

[28] C. Bosbach et al., “Chemical synthesis of site-selective advanced glycation end products in α-synuclein and its fragments,” Organic & Biomolecular Chemistry, vol. 22, no. 13, pp. 2670–2676, 2024, doi: 10.1039/D4OB00225C.

[29] A. B. Uceda, L. Mariño, and M. Adrover, “The Janus face of N-terminal lysines in α-synuclein,” Neural Regeneration Research, vol. 15, no. 10, p. 1840, Oct. 2020, doi: 10.4103/1673-5374.280309.

[30] C. Sun et al., “Cryo-EM structure of amyloid fibril formed by α-synuclein hereditary A53E mutation reveals a distinct protofilament interface,” J Biol Chem, vol. 299, no. 4, p. 104566, Mar. 2023, doi: 10.1016/j.jbc.2023.104566.

[31] “Cryo-EM of full-length α-synuclein reveals fibril polymorphs with a common structural kernel | Nature Communications.” Accessed: Jun. 03, 2025. [Online]. Available: https://www.nature.com/articles/s41467-018-05971-2

[32] K. A. Ehgoetz Martens, C. G. Ellard, and Q. J. Almeida, “Does Anxiety Cause Freezing of Gait in Parkinson’s Disease?,” PLoS ONE, vol. 9, no. 9, p. e106561, Sep. 2014, doi: 10.1371/journal.pone.0106561.

[33] H. C. Roberts, H. E. Syddall, J. W. Butchart, E. L. Stack, C. Cooper, and A. A. Sayer, “The Association of Grip Strength With Severity and Duration of Parkinson’s: A Cross-Sectional Study,” Neurorehabil Neural Repair, vol. 29, no. 9, pp. 889–896, Oct. 2015, doi: 10.1177/1545968315570324.

[34] E. M. Khedr, A. A. Abdelrahman, Y. Elserogy, A. F. Zaki, and A. Gamea, “Depression and anxiety among patients with Parkinson’s disease: frequency, risk factors, and impact on quality of life,” Egypt J Neurol Psychiatry Neurosurg, vol. 56, no. 1, p. 116, Dec. 2020, doi: 10.1186/s41983-020-00253-5.

[35] J. L. Cummings, C. Henchcliffe, S. Schaier, T. Simuni, A. Waxman, and P. Kemp, “The role of dopaminergic imaging in patients with symptoms of dopaminergic system neurodegeneration,” Brain, vol. 134, no. Pt 11, pp. 3146–3166, Nov. 2011, doi: 10.1093/brain/awr177.

[36] O. Stockmann, L. Ye, S. Greten, D. Chemodanow, F. Wegner, and M. Klietz, “Impact of diabetes mellitus type two on incidence and progression of Parkinson’s disease: a systematic review of longitudinal patient cohorts,” J Neural Transm, Jan. 2025, doi: 10.1007/s00702-025-02882-7.

[37] D. Athauda et al., “The Impact of Type 2 Diabetes in Parkinson’s Disease,” Movement Disorders, vol. 37, no. 8, pp. 1612–1623, 2022, doi: 10.1002/mds.29122.

[38] M. B. Thomsen et al., “PET imaging reveals early and progressive dopaminergic deficits after intra-striatal injection of preformed alpha-synuclein fibrils in rats,” Neurobiology of Disease, vol. 149, p. 105229, Feb. 2021, doi: 10.1016/j.nbd.2020.105229.

[39] C. McKinnon et al., “Early-onset impairment of the ubiquitin-proteasome system in dopaminergic neurons caused by α-synuclein,” acta neuropathol commun, vol. 8, no. 1, p. 17, Feb. 2020, doi: 10.1186/s40478-020-0894-0.

[40] F. Weng and L. He, “Disrupted ubiquitin proteasome system underlying tau accumulation in Alzheimer’s disease,” Neurobiology of Aging, vol. 99, pp. 79–85, Mar. 2021, doi: 10.1016/j.neurobiolaging.2020.11.015.

[41] N. Wang et al., “Glial Cell Crosstalk in Parkinson’s Disease: Mechanisms, Implications, and Therapeutic Strategies,” Fundamental Research, Jan. 2025, doi: 10.1016/j.fmre.2024.12.023.

[42] H. S. Kwon and S.-H. Koh, “Neuroinflammation in neurodegenerative disorders: the roles of microglia and astrocytes,” Translational Neurodegeneration, vol. 9, no. 1, p. 42, Nov. 2020, doi: 10.1186/s40035-020-00221-2.

[43] M. Khalid, G. Petroianu, and A. Adem, “Advanced Glycation End Products and Diabetes Mellitus: Mechanisms and Perspectives,” Biomolecules, vol. 12, no. 4, p. 542, Apr. 2022, doi: 10.3390/biom12040542.

[44] D. Indyk, A. Bronowicka-Szydełko, A. Gamian, and A. Kuzan, “Advanced glycation end products and their receptors in serum of patients with type 2 diabetes,” Sci Rep, vol. 11, no. 1, p. 13264, Jun. 2021, doi: 10.1038/s41598-021-92630-0.

[45] J. Tobon-Velasco, E. Cuevas, and M. Torres-Ramos, “Receptor for AGEs (RAGE) as mediator of NF-kB pathway activation in neuroinflammation and oxidative stress,” CNS & neurological disorders drug targets, vol. 13, no. 9, pp. 1615–1626, Dec. 2014, doi: 10.2174/1871527313666140806144831.

[46] V. Deepu, V. Rai, and D. K. Agrawal, “Quantitative Assessment of Intracellular Effectors and Cellular Response in RAGE Activation,” Arch Intern Med Res, vol. 7, no. 2, pp. 80–103, 2024, doi: 10.26502/aimr.0168.

[47] W. Li et al., “Roles of the Receptor for Advanced Glycation End Products and Its Ligands in the Pathogenesis of Alzheimer’s Disease,” Int J Mol Sci, vol. 26, no. 1, p. 403, Jan. 2025, doi: 10.3390/ijms26010403.

[48] K. Sathe et al., “S100B is increased in Parkinson’s disease and ablation protects against MPTP-induced toxicity through the RAGE and TNF-α pathway,” Brain, vol. 135, no. 11, pp. 3336–3347, Nov. 2012, doi: 10.1093/BRAIN/AWS250.

[49] B. W. Festoff, R. K. Sajja, P. van Dreden, and L. Cucullo, “HMGB1 and thrombin mediate the blood-brain barrier dysfunction acting as biomarkers of neuroinflammation and progression to neurodegeneration in Alzheimer’s disease,” J Neuroinflammation, vol. 13, no. 1, p. 194, Aug. 2016, doi: 10.1186/s12974-016-0670-z.

[50] E. Dalfó, M. Portero-Otín, V. Ayala, A. Martínez, R. Pamplona, and I. Ferrer, “Evidence of Oxidative Stress in the Neocortex in Incidental Lewy Body Disease,” Journal of Neuropathology & Experimental Neurology, vol. 64, no. 9, pp. 816–830, Sep. 2005, doi: 10.1097/01.jnen.0000179050.54522.5a.

[51] J. Derk, M. MacLean, J. Juranek, and A. M. Schmidt, “The Receptor for Advanced Glycation Endproducts (RAGE) and Mediation of Inflammatory Neurodegeneration,” J Alzheimers Dis Parkinsonism, vol. 8, no. 1, p. 421, 2018, doi: 10.4172/2161-0460.1000421.

[52] H. Long et al., “Interaction of RAGE with α-synuclein fibrils mediates inflammatory response of microglia,” Cell Reports, vol. 40, no. 12, 2022, Accessed: Dec. 31, 2024. [Online]. Available: https://www.cell.com/cell-reports/fulltext/S2211-1247(22)01238-4?

[53] X. Wang et al., “RAGE Silencing Ameliorates Neuroinflammation by Inhibition of p38-NF-κB Signaling Pathway in Mouse Model of Parkinson’s Disease,” Frontiers in Neuroscience, vol. 14, p. 353, Apr. 2020, doi: 10.3389/FNINS.2020.00353/BIBTEX.

[54] D. O. Peixoto et al., “Increased alpha-synuclein and neuroinflammation in the substantia nigra triggered by systemic inflammation are reversed by targeted inhibition of the receptor for advanced glycation end products (RAGE),” Journal of Neurochemistry, vol. 168, no. 8, pp. 1587–1607, Aug. 2024, doi: 10.1111/jnc.15956.

[55] A. S. Mai, X. Deng, and E.-K. Tan, “Epidemiology of early-onset Parkinson disease (EOPD) worldwide: East versus west,” Parkinsonism & Related Disorders, vol. 129, p. 107126, Dec. 2024, doi: 10.1016/j.parkreldis.2024.107126.

[56] L. A. Volpicelli-Daley et al., “Exogenous α-Synuclein Fibrils Induce Lewy Body Pathology Leading to Synaptic Dysfunction and Neuron Death,” Neuron, vol. 72, no. 1, pp. 57–71, Oct. 2011, doi: 10.1016/j.neuron.2011.08.033.

[57] S. K. Subramanya et al., “A Novel Mouse Model of Parkinson’s Disease for Investigating Progressive Pathology and Neuroprotection,” bioRxiv, pp. 2025–02, 2025.

[58] R. Fiker, L. H. Kim, L. A. Molina, T. Chomiak, and P. J. Whelan, “Visual Gait Lab: A user-friendly approach to gait analysis,” Journal of Neuroscience Methods, vol. 341, p. 108775, Jul. 2020, doi: 10.1016/j.jneumeth.2020.108775.

[59] Z. Suo, M. Wu, B. A. Citron, R. E. Palazzo, and B. W. Festoff, “Rapid tau aggregation and delayed hippocampal neuronal death induced by persistent thrombin signaling,” J Biol Chem, vol. 278, no. 39, pp. 37681–37689, Sep. 2003, doi: 10.1074/jbc.M301406200.

[60] P. Thakur et al., “Modeling Parkinson’s disease pathology by combination of fibril seeds and α-synuclein overexpression in the rat brain,” Proceedings of the National Academy of Sciences of the United States of America, vol. 114, no. 39, pp. E8284–E8293, 2017, doi: 10.1073/pnas.1710442114.

[61] K. Young and H. Morrison, “Quantifying Microglia Morphology from Photomicrographs of Immunohistochemistry Prepared Tissue Using ImageJ,” J Vis Exp, no. 136, p. 57648, Jun. 2018, doi: 10.3791/57648.

[62] S. Bido et al., “Microglia-specific overexpression of α-synuclein leads to severe dopaminergic neurodegeneration by phagocytic exhaustion and oxidative toxicity,” Nat Commun, vol. 12, no. 1, p. 6237, Oct. 2021, doi: 10.1038/s41467-021-26519-x.

