## Supplementary material for "Glycated alpha-synuclein assemblies cause distinct Parkinson’s disease pathogenesis in mice"

Supplementary information containing figures S1- S5

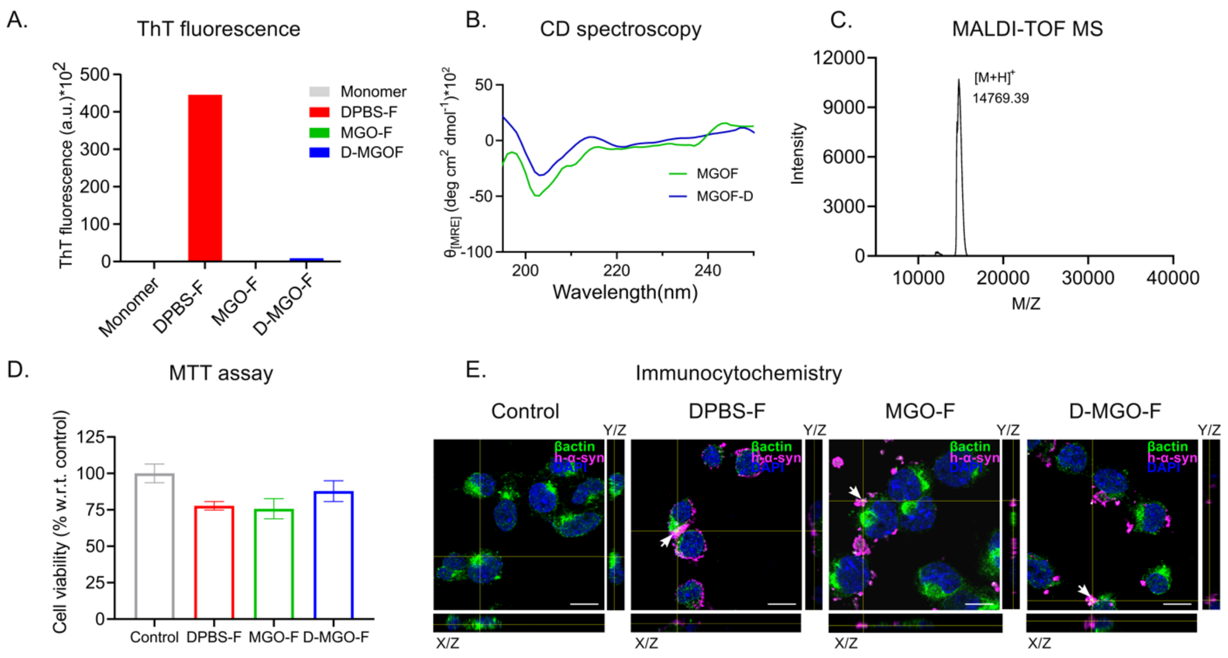


**Figure S1- Dialyzed MGO-F showed similar characteristics to non-dialyzed MGO-F.**

A. Representative ThT fluorescence of recombinant α-Syn 96-hours post-incubation in fibrillazation conditions, with or without MGO and dialyzed MGO-F (D-MGO-F), shows lack of beta sheet structure in both MGO-F and D-MGO-F compared to DPBS-F. B. CD spectra of MGO-F and D-MGO-F show a similar predominantly random-coiled structure. C. MALDI-TOF MS spectra of D-MGO-F shows similarity to the spectra of MGO-F. D. MTT cell viability assay of N2a cells after 48 hours exposure to DPBS-F, MGO-F and D-MGO-F (n=3); data plotted as mean ± SEM. E. Representative micrographs with orthogonal views showing co-staining for beta-actin, human α-Syn and DAPI (scale bar-10 μm), marked by white arrows.

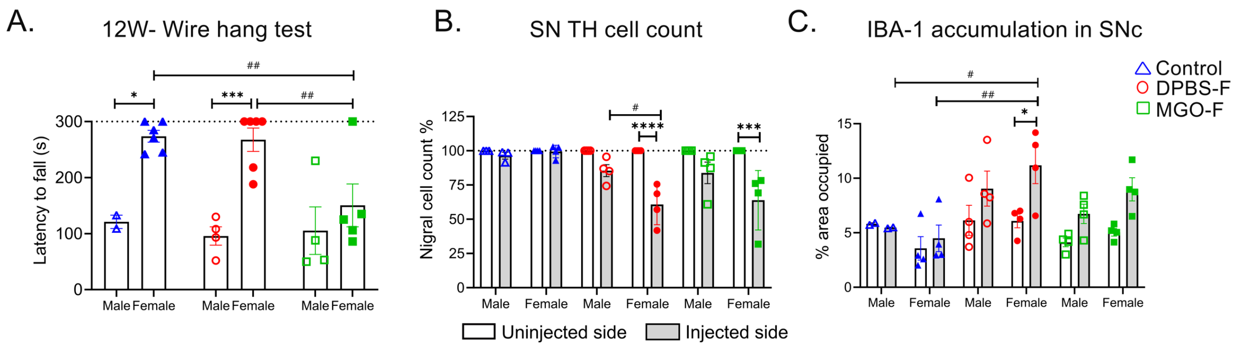


**Figure S2- Sex-based differences in various treatment groups**

Wire hang test 12 weeks post-surgery shows MGO-F female mice have a greater loss in latency to fall than female mice of other groups. Males in the control and DPBS-F groups performed worse compared to females in their group. B. Quantification of TH-positive cells in SN shows higher neurodegeneration in female mice of the DPBS-F and MGO-F groups. C. Quantification of the percentage area occupied by IBA-1 in the SNc region, showing higher inflammation in the injected side of female mice of the DPBS-F and MGO-F groups. All data represented as mean ± SEM, n=2-6 per sex; Mixed two-way ANOVA followed by Tukey's or Sidak's multiple comparison test; *$p < 0.05$, ***$p < 0.001$, $^{\#}p < 0.05$, $^{\#\#}p < 0.01$; * indicates comparison within groups, and # represents comparison between groups; Filled symbols-female and empty symbols-male.

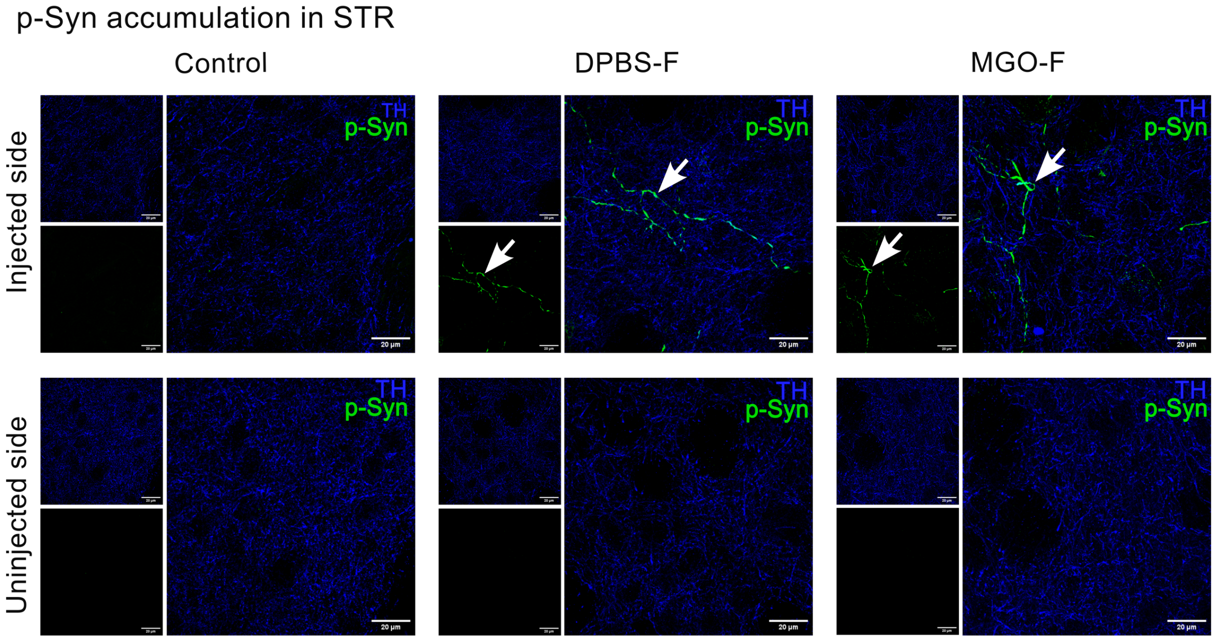


**Figure S3- Accumulation of p-Syn in the striatum**

Representative micrographs of STR co-stained with p-Syn and TH in all three indicated groups (scale bar-20 µm). Arrows point to the p-Syn deposits.

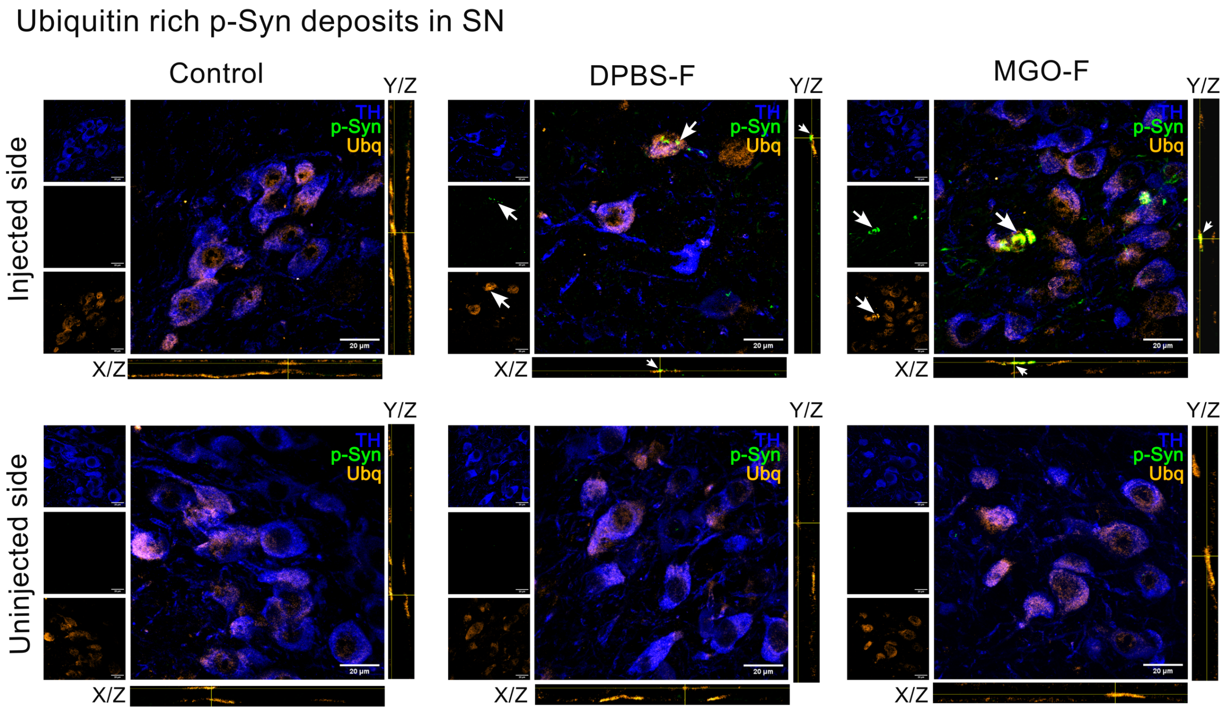


**Figure S4- Ubiquitin-accumulation in the SN**

Representative micrographs of SN co-stained with TH, p-Syn and Ubq (scale bar-20 µm) with orthogonal views. Arrows point to the p-Syn aggregates showing colocalization with ubiquitin.

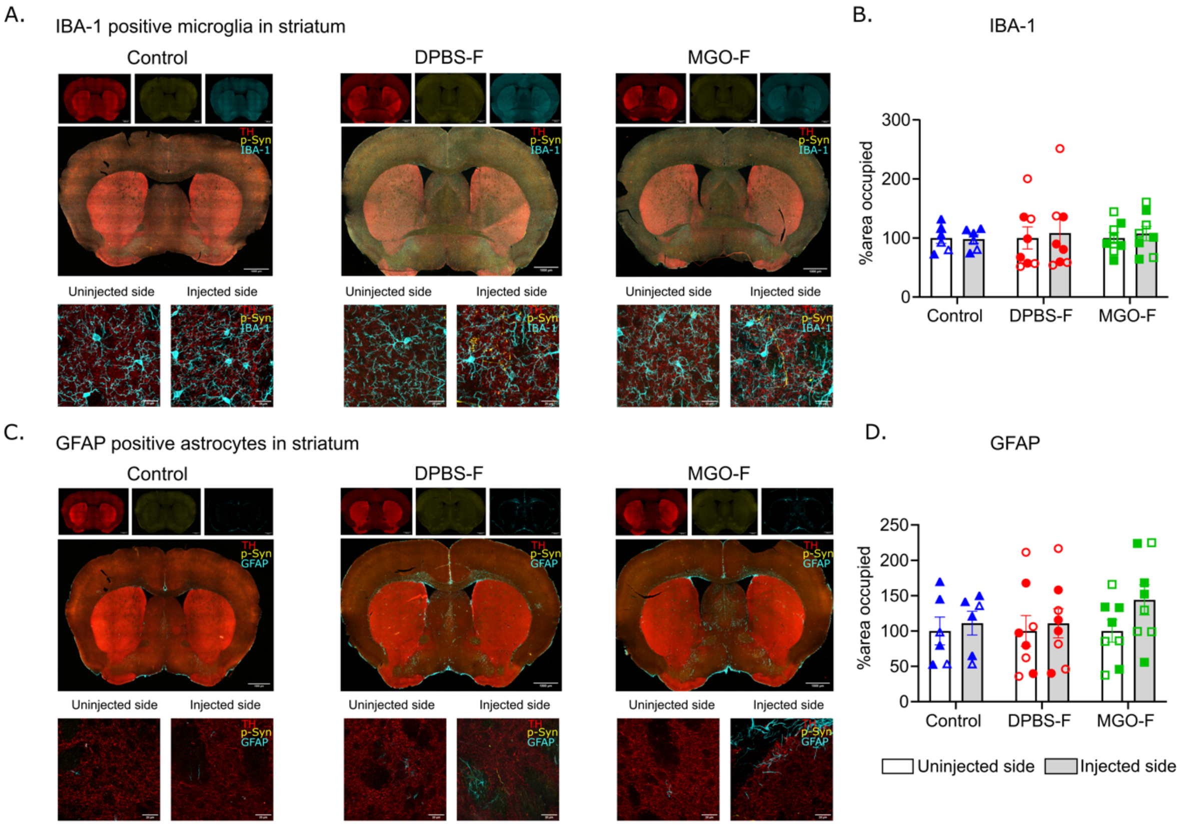


**Figure S5- Neuroinflammation in striatum.**

A. Representative micrographs of STR co-stained for TH, p-Syn and IBA-1 (scale bar-1000 μm) and their representative higher magnification images below (scale bar 20-μm). B. Quantification of the percentage area occupied by IBA-1 in STR shows no change across different groups. C. Representative micrographs of STR co-stained for TH, pSyn and GFAP (scale bar-1000 μm) and their representative higher magnification images below (scale bar-20 μm). D. Quantification of the percentage area occupied by GFAP in STR. Data represented are as mean ± SEM; n=6-8 mice for all groups; Mixed two-way ANOVA followed by Tukey's or Sidak's multiple comparison test; Filled symbols- female and empty symbols- male.
